# Addressing the pooled amplification paradox with unique molecular identifiers in single-cell RNA-seq

**DOI:** 10.1101/2020.07.06.188003

**Authors:** Johan Gustafsson, Jonathan Robinson, Jens Nielsen, Lior Pachter

## Abstract

The incorporation of unique molecular identifiers (UMIs) in single-cell RNA-seq assays allows for the removal of amplification bias in the estimation of gene abundances. We show that UMIs can also be used to address a problem resulting from incomplete sequencing of amplified molecules in sequencing libraries that can lead to bias in gene abundance estimates. Our method, called BUTTERFLY, is based on a zero truncated negative binomial estimator and is implemented in the kallisto bustools single-cell RNA-seq workflow. We demonstrate its efficacy using a range of datasets and show that it can invert the relative abundance of certain genes in cases of a pooled amplification paradox.

## Background

Droplet-based single-cell RNA sequencing (scRNA-Seq) technologies have made possible quantification of transcriptomes in individual cells at a large scale (1), enabling the study of the diversity in molecular state among cells. As an increasing number of datasets have been collected (2), methods for integration of results derived by different laboratories have become paramount. The diversity of experimental and computational methods used in producing individual datasets makes careful accounting of technical and batch effects essential (3).

Most single-cell RNA-seq technologies require amplification of the RNA starting material via PCR, a step that is known to introduce bias across genes depending on the nucleotide sequence. For example, amplification has been shown to depend on the GC content of a gene (4). Single-cell RNA-Seq can require many PCR amplification cycles, even more than bulk sequencing, due to the small amount of mRNA molecules available in each cell. Fortunately, the introduction of unique molecular identifiers (UMIs) (5), where all mRNA molecules are tagged with random barcodes, can be used to account for PCR duplicates, since copies of captured mRNA molecules can be detected and discarded. This process is known as deduplication, or UMI collapsing (6). However, UMI collapsing does not address another bias that can result from incomplete representation of differentially amplified molecules in sequenced products from a library (Figure 1A), and is common in droplet-based single-cell RNA-Seq experiments (7).

**Figure 1:**
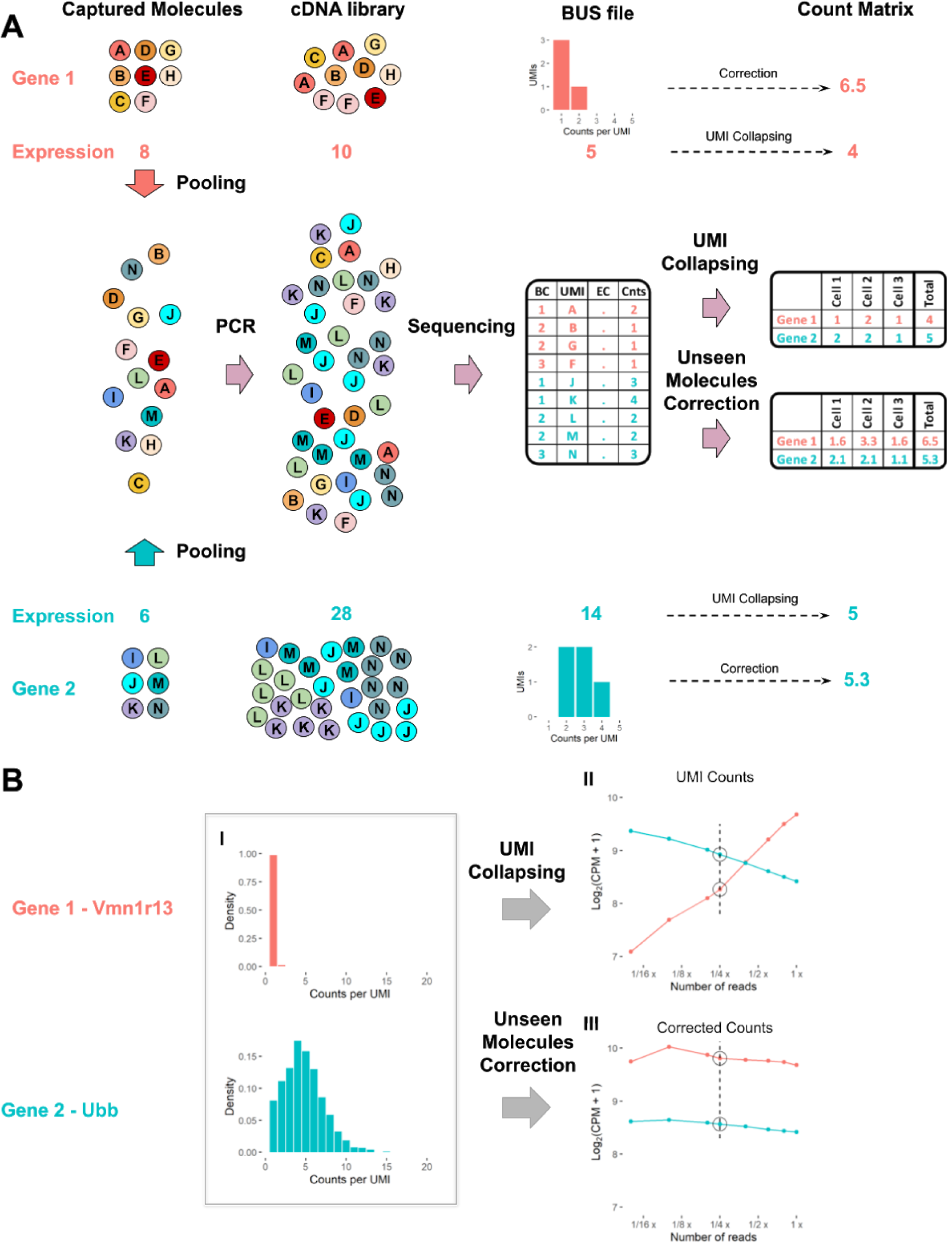
The pooled amplification paradox. A. Illustration of a seemingly paradoxical reversal in gene abundance estimates arising from incomplete sampling of a cDNA library generated after a differential amplification of two genes. Consider a situation where more molecules have been captured for Gene 1 (8) than Gene 2 (6). Despite the correction for amplification bias with unique molecular identifiers, the increased amplification of Gene 2 relative to Gene 1, coupled with the incomplete sequencing of the cDNA library, results in a lower copy number estimate of Gene 1 (4) in comparison to Gene 2 (5). Correction of abundance estimates due to unseen molecules using a histogram of counts per UMI improves the quantification, reversing the relative amount of Gene 1 and Gene 2. B. Example of the pooled amplification paradox as seen in an analysis of the EVAL dataset. I) Histograms of the counts per UMI for two genes with different amplification. The histograms are generated from downsampled data (4 times), visualized by a dashed line in II and III. II) Change in the mean gene expression estimates across cells of the two genes when downsampling the reads. III) Change in gene expression for downsampled data when correcting for unseen molecules. The code to reproduce this figure is here: <u>code for Figure 1</u>

Estimation of unseen species (8,9) is commonly used in ecology, where the number of species encountered in an ecosystem given a certain sampling effort is estimated from a limited number of samples. We translate this to the problem of estimating the unseen number of molecules in a single-cell sequencing experiment, where the species correspond to unique molecules and samples to sequencing reads. We show that this is possible in assays that utilize unique molecular identifiers.

The mathematical problem of estimating unseen species has previously been addressed with the introduction of several estimators. The simplest is the Good-Toulmin estimator (8), which while easy to utilize, is limited in prediction range, producing stable predictions of species represented only in up to twice the number of samples. Although this limitation can be addressed (10), the estimator is still not practically useful for sequencing experiments. Daley et al. therefore developed Preseq (11–13), which does not suffer from such limitations, and employed it to estimate the optimal number of reads for genome sequencing. Their method is based on rational function approximation, which substantially increases the radius of convergence of the power series appearing in the Good-Toulmin estimator, providing a stable estimator.

Here, we present BUTTERFLY, a method that utilizes estimation of unseen species for addressing the bias caused by incomplete sampling of differentially amplified molecules. Specifically, we extrapolate the gene expression to a higher number of reads by estimating the missing number of molecules per gene, validate the results using two different approaches, and demonstrate that BUTTERFLY can be efficiently applied to datasets using the kallisto bustools single-cell RNA-seq workflow (14).

## Results

To measure the extent and heterogeneity of amplification bias we analyzed a total of 13 scRNA-seq datasets which were generated from a total of 5 different technologies (see Methods). We found that the variation in RNA amplification was substantial across genes (Supplementary Figures 1 -- 3). Amplification bias patterns were different across technologies. For example, all 10X technologies (Supplementary Figure 1) displayed consistently more amplification bias than Drop-seq (Supplementary Figure 2) or Seq-Well (Supplementary Figure 3). Furthermore, we found that amplification bias is consistent across technologies. For example, the variance in the fraction of single copy molecules (FSCM) in 10X technologies was consistently 0.024 (std. dev. 0.008) whereas the variance in FSCM in the Drop-seq datasets was 0.013 (std. dev. 0.004). We also found that specific genes were more likely to be affected by amplification bias than others (Figure 2A, Supplementary Figure 4). Interestingly, the genes with high amplification bias, as determined by their FSCM, are generally the same across datasets, especially in data generated with the same technology (Figure 2B, Supplementary Figure 5).

**Figure 2:**
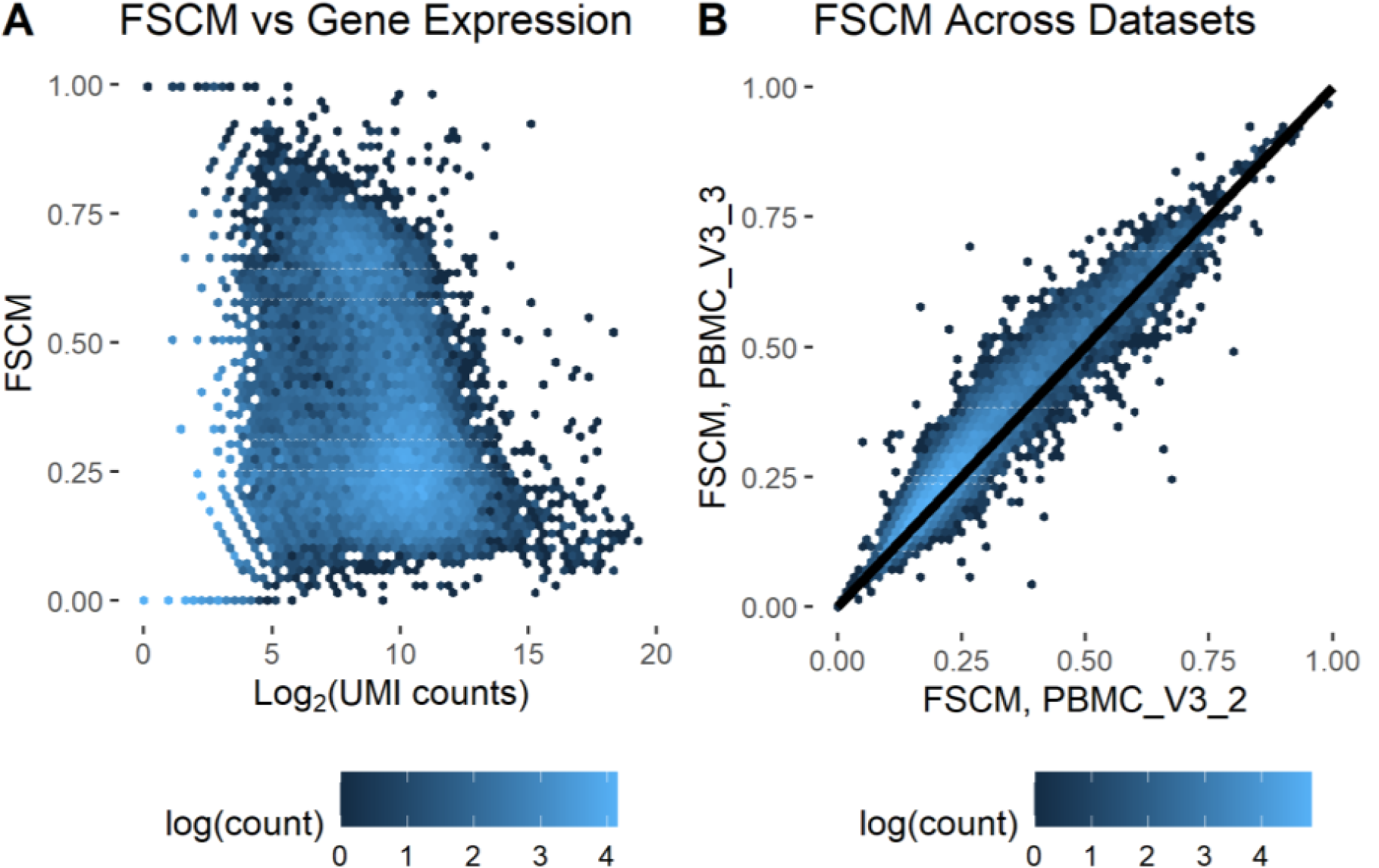
Differences in fraction of single-copy molecules across genes. A. Fraction of single copy molecules (FSCM) vs mean gene expression across cells for the PBMC_V3_3 dataset. B. Comparison of gene FSCM between 2 datasets (PBMC_V3_2 and PBMC_V3_3) produced using the same technology (10x Chromium v3). The code to reproduce this figure is here: <u>code for Figure 2</u>

**Figure 3:**
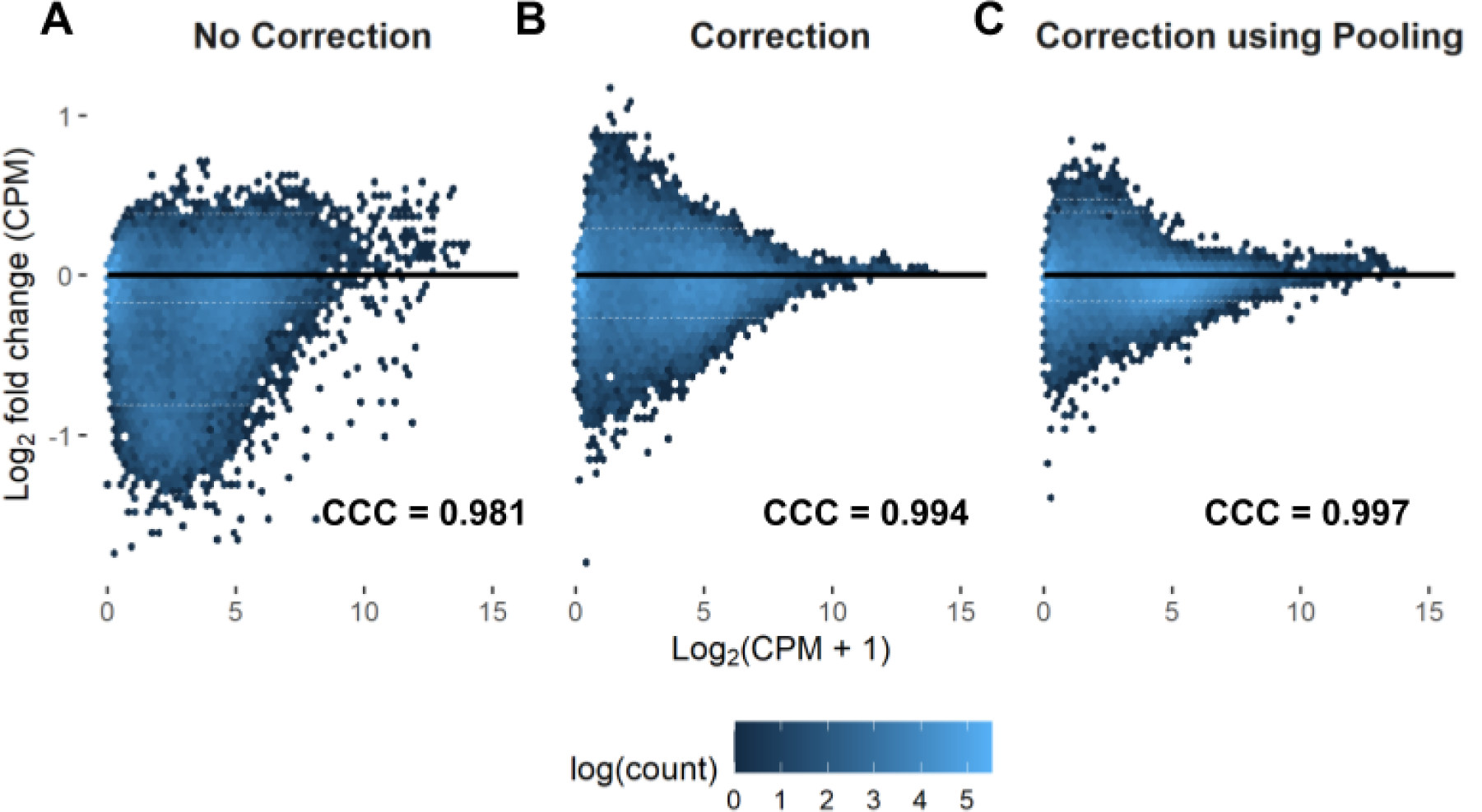
Effect of BUTTERFLY correction on gene expression. The log fold change in gene expression for the PBMC_V3_3 dataset between the downsampled data (at 1/10 number of reads) and the original data. A. No unseen molecules correction. B. Unseen molecules correction applied to every gene based on the data within the dataset. C. The same correction as in B, but using additional data from six other 10x Chromium datasets for estimating the copies per UMI histograms per gene (Methods). The code to reproduce this figure is here: <u>c ode for Figure 3</u>

**Fig. 4:**
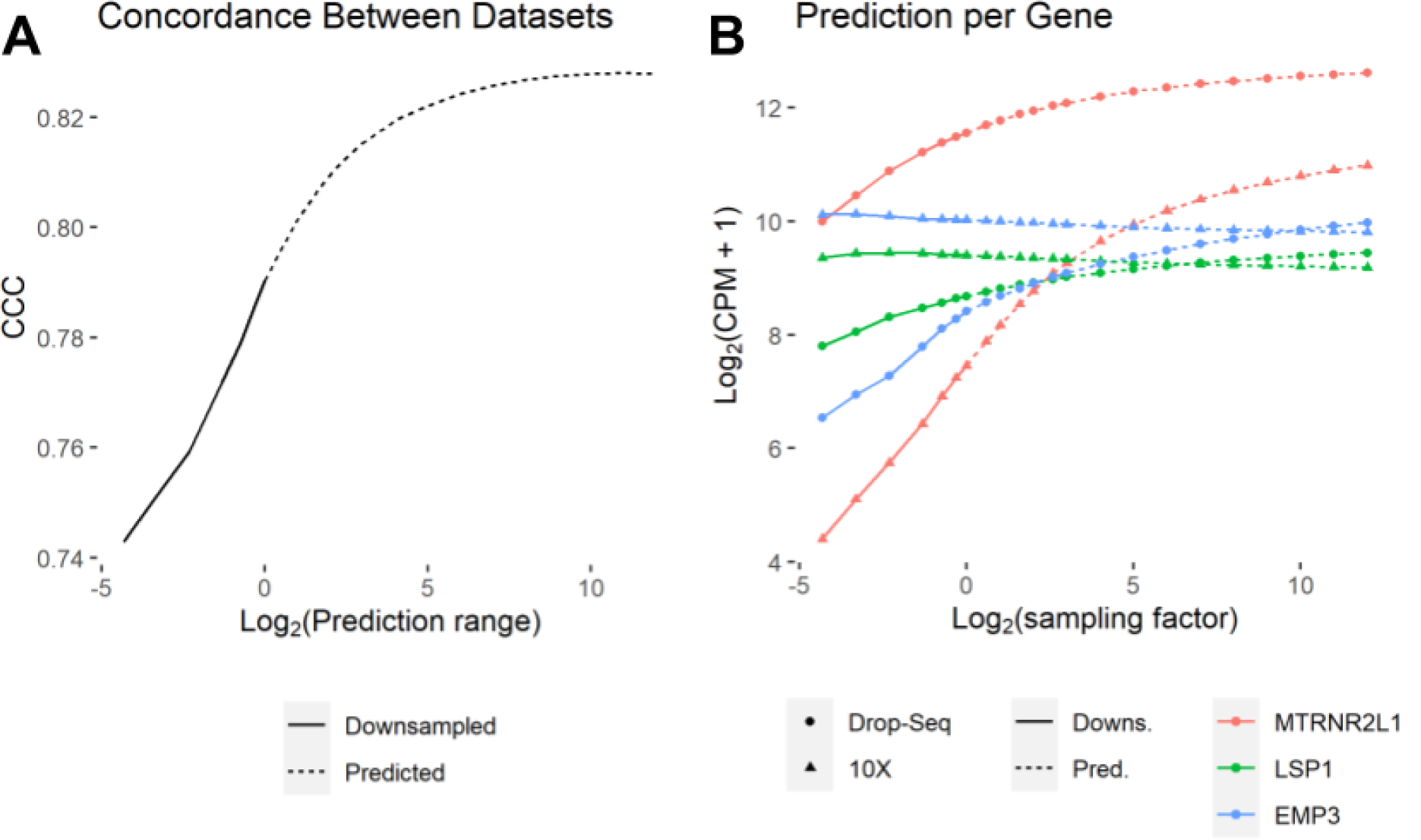
Unseen molecules correction increases gene expression similarity across technologies. Two PBMC datasets generated with different technologies ((EVALPBMC, 10x Chromium v2, and EVALPBMC_DS, Drop-Seq) were downsampled to varying depths, and corrected for unseen molecules using BUTTERFLY. A. The figure shows the change in correlation (CCC) between the two datasets at different prediction/downsampling points. For the red curve, the 10X dataset is predicted/downsampled as defined by the prediction range, while the Drop-Seq dataset is kept constant at the full number of reads (no prediction). For the blue line, both datasets are predicted/downsampled at each point. B. Change in gene expression from prediction/downsampling for three genes, for both datasets. The code to reproduce this figure is here: <u>code for Figure 4</u>

To assess the implications of amplification bias on single-cell RNA-seq quantification, we downsampled reads in a mouse 10X brain cortex dataset (the EVAL dataset, see Methods) and analyzed quantification of abundances with standard UMI collapsing for a pair of genes (Figure 1B). We found that while the vomeronal 1 receptor 13 gene (*Vmn1r13*) appeared to be 2.4 times more highly expressed than the Ubiquitin B gene (*Ubb*) in the full dataset (30 M reads), when downsampling to 1.5 M reads (1/20th the dataset) the opposite was the case: *UBB* was 4.9 times more highly expressed than *vmn1r13*. Increasing the number of reads yields the discovery of many new molecules for *vmn1r13*, but few for *Ubb*, since many more of the molecules belonging to *Ubb* have already been sampled, and are therefore “canceled out” during UMI collapsing.

We hypothesized that by predicting unseen species (8,9), we could reduce, or eliminate, the bias introduced by naive UMI collapsing. Each gene’s expected increase in gene expression with more reads can be estimated from measurements of counts per UMI (CU histogram, see Figure 1B i). We evaluated several methods for this purpose (Supplementary Figure 6): The Daley et al. (11) Preseq method (Preseq DS) exhibits similar performance to the zero-truncated negative binomial (ZTNB) (9), and both are more stable than the Good-Toulmin estimator (8). We found that ZTNB has better performance for highly expressed genes, while Preseq DS performs slightly better for medium range genes for some datasets (see Supplementary Note).

To evaluate the performance of BUTTERFLY we downsampled a human 10X peripheral blood mononuclear cell dataset (PBMC_V3_3, see Methods) to one tenth of the reads and compared the uncorrected gene expression estimates and BUTTERFLY corrected gene expression estimates in the downsampled dataset, to those of the full dataset. Figure 3A shows the log fold change for all genes where no correction is applied, showing large discrepancies for many genes. The concordance correlation coefficient (CCC) was 0.981 (see Methods). With BUTTERFLY the difference is clearly reduced (Figure 3B, CCC = 0.994), especially for highly expressed genes. These results are recapitulated in 12 other datasets from a variety of technologies including 10X Genomics v2, v3 and Next Gem, Drop-seq and SeqWell (Supplementary Figures 7 -- 19). We found that pooling histograms derived from distinct datasets further improves results; we assembled CU histogram data from 6 other 10X Chromium datasets from human PBMC and utilized it for prediction (Methods). Use of pooled data clearly improves the prediction for low expression genes (Fig. 4C, CCC = 0.997).

Part of the discrepancy between corrected expression of downsampled data and the full datasets can be explained by sampling effects. To estimate the magnitude of that effect, we repeatedly downsampled the original dataset to one tenth of the reads for 20 iterations and calculated the mean expression for each gene, thereby obtaining a bound on the accuracy possible (Supplementary Figure 20).

While the downsampling results show that BUTTERFLY correction can be used to scale the gene expression of each gene to resemble the gene expression that more reads would yield, they do not necessarily imply that the corrected expression values are closer to ground truth. To investigate whether that is indeed the case, we assessed whether BUTTERFLY increases similarity of gene abundance estimates between datasets from the same biological sample assayed with different technologies. Figure 4A shows that as a correction for unseen molecules is applied to both 10X Chromium and Drop-seq datasets from PBMC cells, there is an increase in concordance. Figure 4B shows that the improvement in concordance is driven by individual genes, as the effect on concordance resulting from correction of unseen molecules on individual genes is highly gene dependent.

## Discussion

The “pooled amplification” paradox we have highlighted is of significant concern because it can affect some genes significantly, and therefore inferences based on their relative abundance estimates may be based on biased information. For example, the gene NEUROD1 has a very high fraction of single-copy molecules in 10x Chromium datasets (FSCM = 0.90, calculated from the joint data of all 10x PBMC datasets used in this study) suggesting that the number of cells with a detected expression of this gene is much lower than what would otherwise be expected. This gene has for example been identified as a marker gene used to identify enteroendocrine cell precursors (15). Our results show that this gene may appear to be underrepresented in data due to low PCR amplification.

In addition to improving abundance estimates of specific genes, we have shown that BUTTERFLY can help reduce batch effect between datasets sequenced at different depths. This result should be interesting to explore in conjunction with single-cell integration methods (16). Furthermore, while we have demonstrated BUTTERFLY in the context of single-cell RNA-seq, the approach we have outlined is relevant for any assay in which objects are sampled after amplification, and where the pooled amplification paradox may occur (see, e.g. (17)). In particular, genomics assays utilizing UMIs should benefit from our method. There are also other applications of estimation of unseen species in genomics, as recently shown in (18).

Our work also highlights the need for reliable estimators for the unseen species problem in the case where count histograms are based on few observations. While there has been significant effort expended in the development of estimators in the case of abundant data, this setting may be ripe for new discoveries. Furthermore, the improved reliability of estimators with more data means our approach may induce bias in accuracy favoring highly abundant genes. This should be taken into account in downstream analyses and in interpretation of results, much in the same way that improved abundance estimates of long genes affect interpretation of differential analysis (19).

## Methods

### Datasets

We analyzed a total of 13 public scRNA-Seq datasets collected using 5 technologies; Drop-Seq, Seq-Well and 10x Chromium version v2, v3, and Next GEM. The datasets were generated from 4 sample types; mouse brain (dataset EVAL), mouse retina (datasets MRET and MRET2), human lung tumor (dataset LC), and human PBMC (remaining datasets). The datasets, including metadata, are listed in Table S1, with additional information in Supplementary Table S2. The datasets were processed using kallisto (20) and bustools (14), yielding counts and gene association for each unique molecule. Molecules mapped to multiple genes were discarded before analysis.

### Pre-processing of Sequencing Files

The datasets were processed using kallisto (20) version 0.46.2 and and a version of bustools (14) specifically developed for this study, yielding counts and gene association for each unique molecule. kallisto index files were created for mouse and human using cDNA files from Ensembl (v. 96 for mouse, v. 94 for human).

We developed the new commands *collapse* and *umicorrect* in a branch of the bustools code, and modified the commands text and fromtext, to enable production of bus files mapped per gene (BUG-files) for further processing in R. The umicorrect command was implemented as described previously(21), while collapse transforms the BUS records from being mapped to equivalence classes to being merged where appropriate and mapped to genes. Transcripts to genes files, which are required for the collapse command, were generated with the function transcript2gene from the BUSpaRse R package(22), version 0.99.25.

### Processing of data in R

All reads mapping to more than one gene were discarded before further processing, as were cells with fewer than 200 UMIs (except for the LC dataset, where cells with fewer than 1000 UMIs were discarded, motivated by the large number of cells compared to the expected number from the authors). Statistic metrics were also calculated for each dataset (Supplementary Table S2).

### Correction for unseen molecules

To correct gene abundance estimates, the number of unseen molecules are predicted for each gene, assuming that the same gene behaves similarly across cells. For each gene, all molecules across all cells are pooled and used to calculate a CU histogram. The CU histogram is then in turn used as input to a prediction algorithm. Prediction was done using the Good-Toulmin estimator as well as the DS (based on rational functions approximation) and zero truncated negative binomial (ZTNB) included in the PreseqR package. We implemented the Good-Toulmin estimator in R and used the functions ds.rSAC and a modified version of ztnb.rSAC from PreseqR (where the modified version has larger error tolerances, which speeds up computation time considerably while producing very similar results). The gene expression was predicted per gene, pooling the UMIs from all cells in the dataset. Histograms over the number of copies per UMI (CU histograms) were constructed per gene and used as input to the prediction algorithms. The predicted number of UMIs per gene was then used to calculate the gene expression in counts per million (CPM). We used ZTNB for prediction except for figure 3C, where the pooled prediction is based on the DS algorithm (MT = 2).

The ZTNB method is based on using an expectation-maximization algorithm for fitting a negative binomial curve to the histogram of number of copies per UMI, where the number of molecules with zero counts is unknown. It is then assumed that the size parameter of the negative binomial remains the same as the number of reads increase, and that the mean is proportional to the number of reads. The number of molecules with at least one copy can then be estimated from the probability density function of the negative binomial.

Once the predicted gene expression is estimated, the UMI counts in the counts matrix are scaled by a factor *m*_*g*_, calculated for each gene *g* as

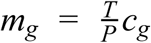

where *T* is the total number of UMIs in the count matrix, *P* is the total number of predicted UMIs across all genes, and *c*_*g*_ is the number of predicted UMIs for gene *g*.

### Correction using Pooled Data from Other Datasets

The prediction is dependent on having enough data per gene to build CU histograms, as sampling effects on the histogram leads to unstable prediction. A way to circumvent this issue is to use CU histogram data from other similar datasets in the prediction. The Preseq DS algorithm with CU histograms truncated at 2 copies per molecule is convenient in this regard, since we only need to estimate the fractions of molecules that have one (FSCM) and two (FDCM) copies. These metrics were measured per gene for the datasets PBMC_V3, PBMC_V3_2, PBMC_NG, PBMC_NG_2, PBMC_V2 and EVALPBMC (pool source datasets), and were used to predict the dataset PBMC_V3_3.

Since each dataset has a different degree of saturation (i.e. average counts per UMI), there is a need to normalize FSCM and FDCM between the dataset being predicted and the pool source datasets (see for example Fig. 3C, where the dataset PBMC_V2 on average has a lower FSCM than PBMC_V3_3). We utilized quantile normalization to adjust the FSCM and FDCM of the pool source datasets to be more similar that of the dataset being predicted.

The pooled FSCM and FDCM metrics for a gene is then calculated as a weighted mean of all datasets, including the dataset being predicted, where the weight is the number of UMIs for the gene per dataset. A third metric, the fraction of molecules that have more than two copies (FMCM), is calculated from FSCM and FDCM, as FMCM = 1 - FSCM - FDCM. The histogram used for prediction is simplified to these three bins, and constructed using those metrics scaled with the number of UMIs of the gene in the dataset to be predicted. Prediction is then carried out as described above.

For generating Figure 3, where prediction is performed using downsampled data, the pool source datasets were downsampled as well to better match the dataset being predicted regarding degree of saturation. Since much data is lost during downsampling, each dataset was repeatedly downsampled 10 times and added to the data pool, providing a list of in total 60 datasets.

### Concordance Correlation Coefficient

To compare the similarity in gene expression of two samples, the Pearson correlation is not a suitable metric since it measures linearity, and not similarity. We instead used Lin’s concordance correlation coefficient (CCC), which describes the expected perpendicular distance from a 45° line passing through the origin. We used the CCC function in the R package DescTools (version 0.99.36) to calculate the metric, using default parameters.

The similarity was calculated on log-transformed data, where the transformed gene expression *l*_*i*_ for gene *i* is calculated as *l*_*i*_ = *log*_2_(*e*_*i*_ + 1), where *e*_*i*_ is the gene expression of the gene in counts per million (CPM).

### Figure Details

In figure 1 B III, the correction is based on correcting the UMIs for all genes individually. The prediction range is the same for all genes, scaling up the counts to match the total counts in the full dataset. The predicted UMIs are then CPM normalized.

To avoid the uncertainty in the FSCM calculations that arise from having too few molecules, only genes with at least 200 molecules in both datasets were included in figure 2B and supporting figure 5.

To avoid the uncertainty in prediction from lowly expressed genes, genes with a gene expression lower than 100 CPM were removed before calculating CCC in figure 4, leaving in total 1018 genes. CPM was then recalculated based on these genes only to avoid the influence of lowly expressed genes, for which prediction is less accurate.

In all figures where the gene expression of a dataset is used, the gene expression is calculated as the mean gene expression across all cells.

### Implementation

The ZTNB prediction was implemented in bustools with the addition of the ‘predict’ command using the same algorithm as PreSeqR(12,13), and utilizing the c++ libraries Eigen(23), LBFGSpp(24), and CppOptimizationLibrary(25). The count command in bustools has been extended with the option “--hist”, which generates CU histograms that serve as input to predict, together with the count matrix. In addition, UMI correction was implemented as the “umicorrect” command, utilizing the method of (21).

## Declarations

### Availability of data and materials

Means to access the datasets analyzed during the current study are listed in Supplementary table S1. The source code as well as Jupyter notebooks for generating the figures is available at: https://github.com/pachterlab/GRNP_2020. The source code for the branch of bustools used in this project is available at: https://github.com/BUStools/bustools/tree/butterfly/src.

## Acknowledgements

We thank Pall Melsted, Sina Booeshaghi and Joseph Min for helpful suggestions on the project and on the integration of BUTTERFLY in bustools.

## Authors’ Contribution

Conceptualization, J.G., L.P.; Methodology, J.G, L.P.; Software, J.G.; Writing – Original Draft, J.G., L.P.; Writing – Review & Editing, J.G., L.P., J.R., J.N.; Supervision, L.P., J.R., J.N.; Funding Acquisition, L.P., J.R., J.N.

