## Supplementary information for "Addressing the pooled amplification paradox with unique molecular identifiers in single-cell RNA-seq"

* Corresponding author

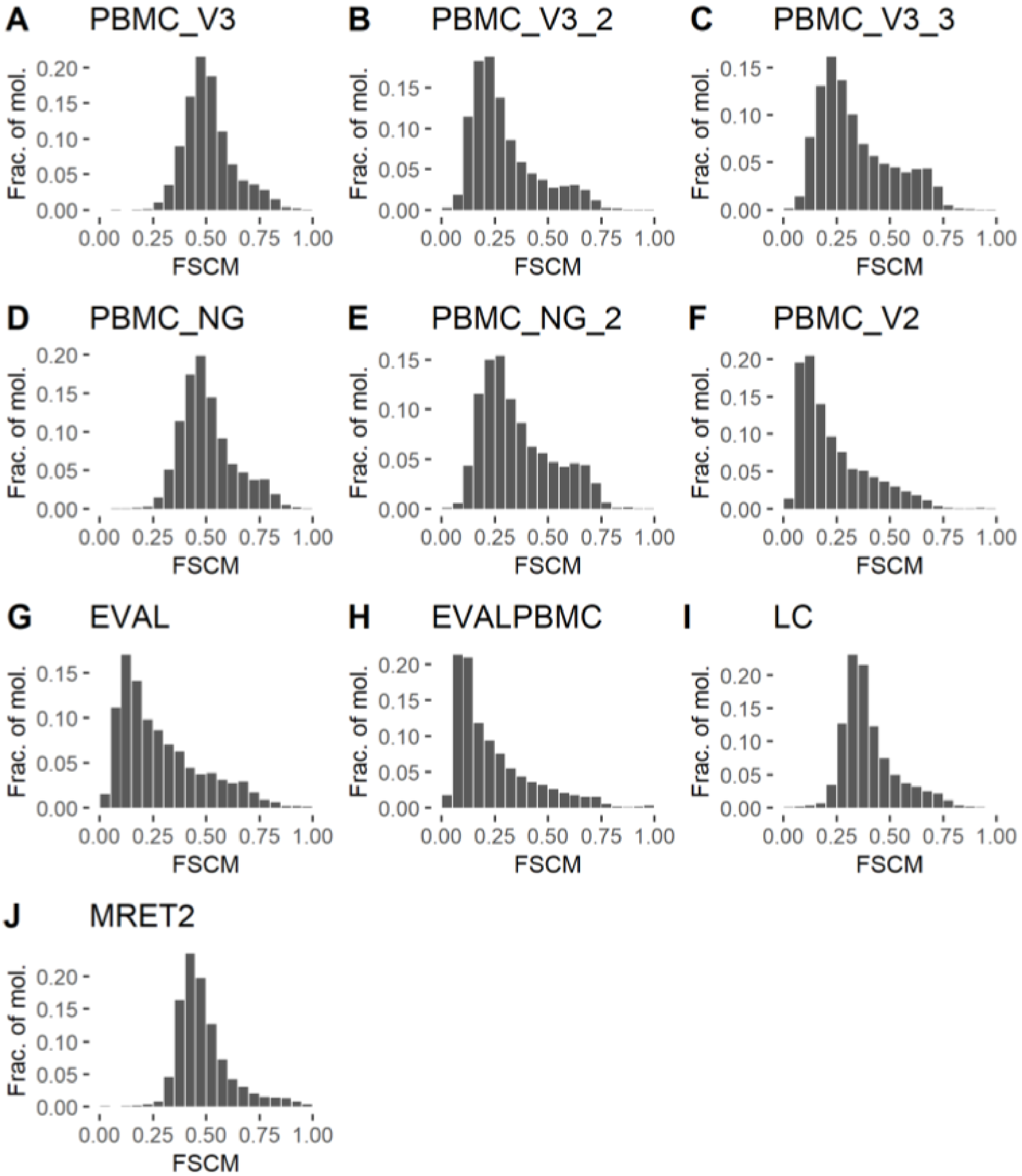


***Fig S1: Histograms over fraction of single-copy molecules per gene for 10x Chromium datasets.*** *Genes with fewer than 200 molecules present in the dataset are not shown. The code to reproduce this figure is here: code*

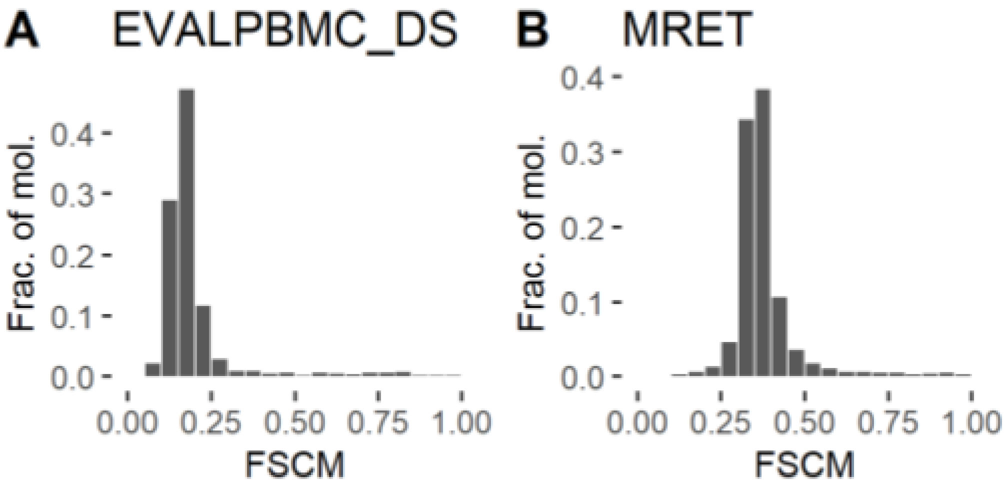


***Fig S2: Histograms over fraction of single-copy molecules per gene for Drop-Seq datasets.*** *Genes with fewer than 200 molecules present in the dataset are not shown. The code to reproduce this figure is here: code*

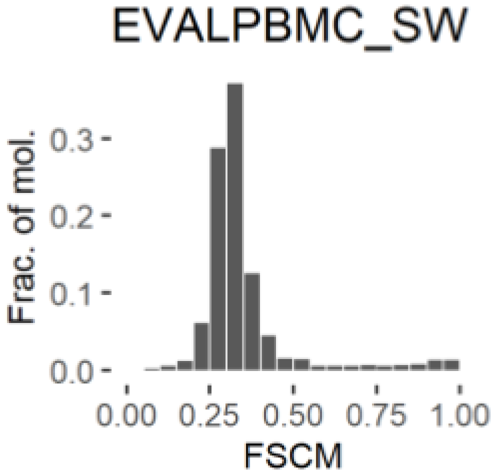


***Fig S3: Histogram over fraction of single-copy molecules per gene for the Seq-Well dataset.*** *Genes with fewer than 200 molecules present in the dataset are not shown. The code to reproduce this figure is here: [code]*

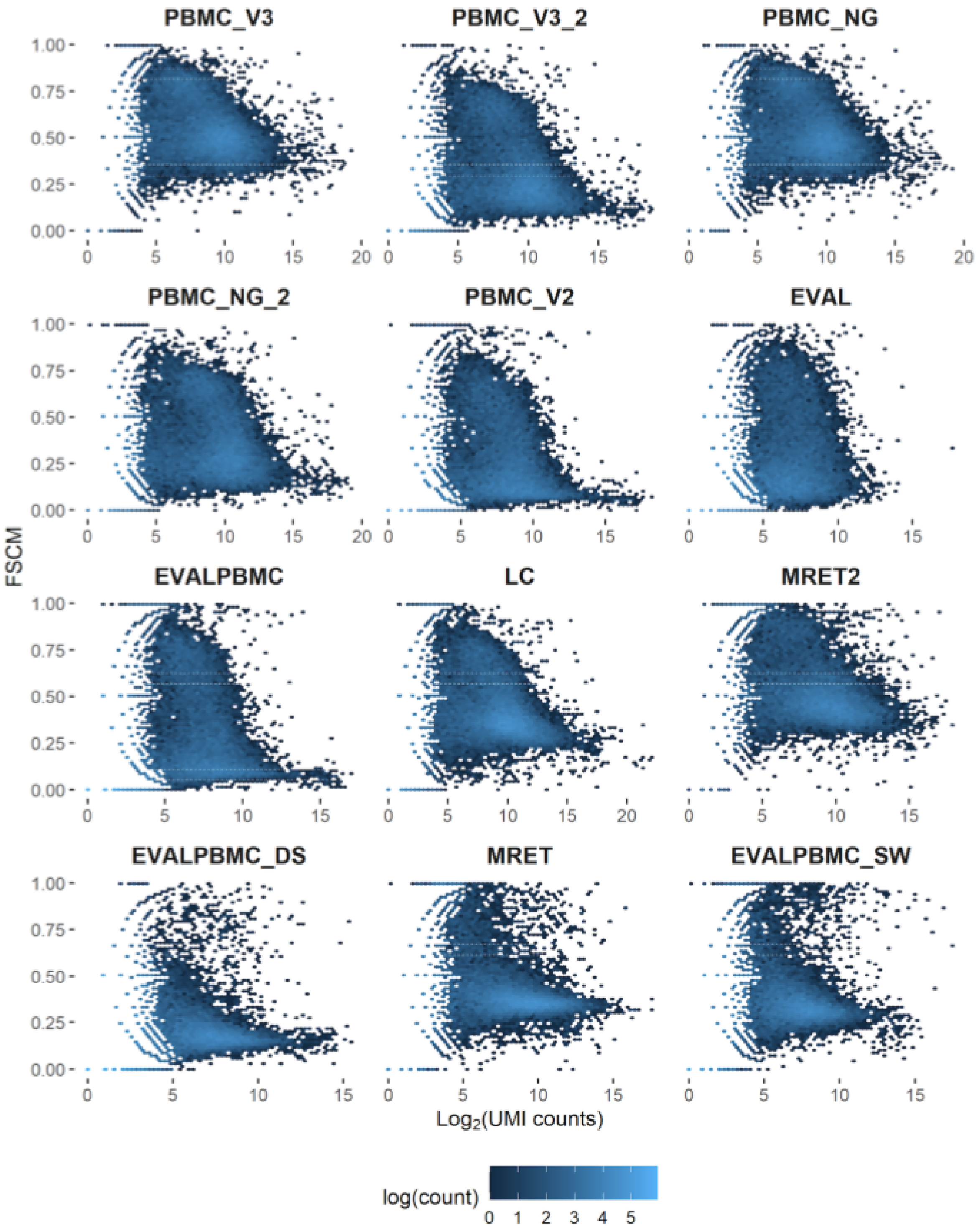


***Fig. S4: Fraction of single-copy molecules vs number of UMIs per gene per dataset.***
*The code to reproduce this figure is here: [code]*

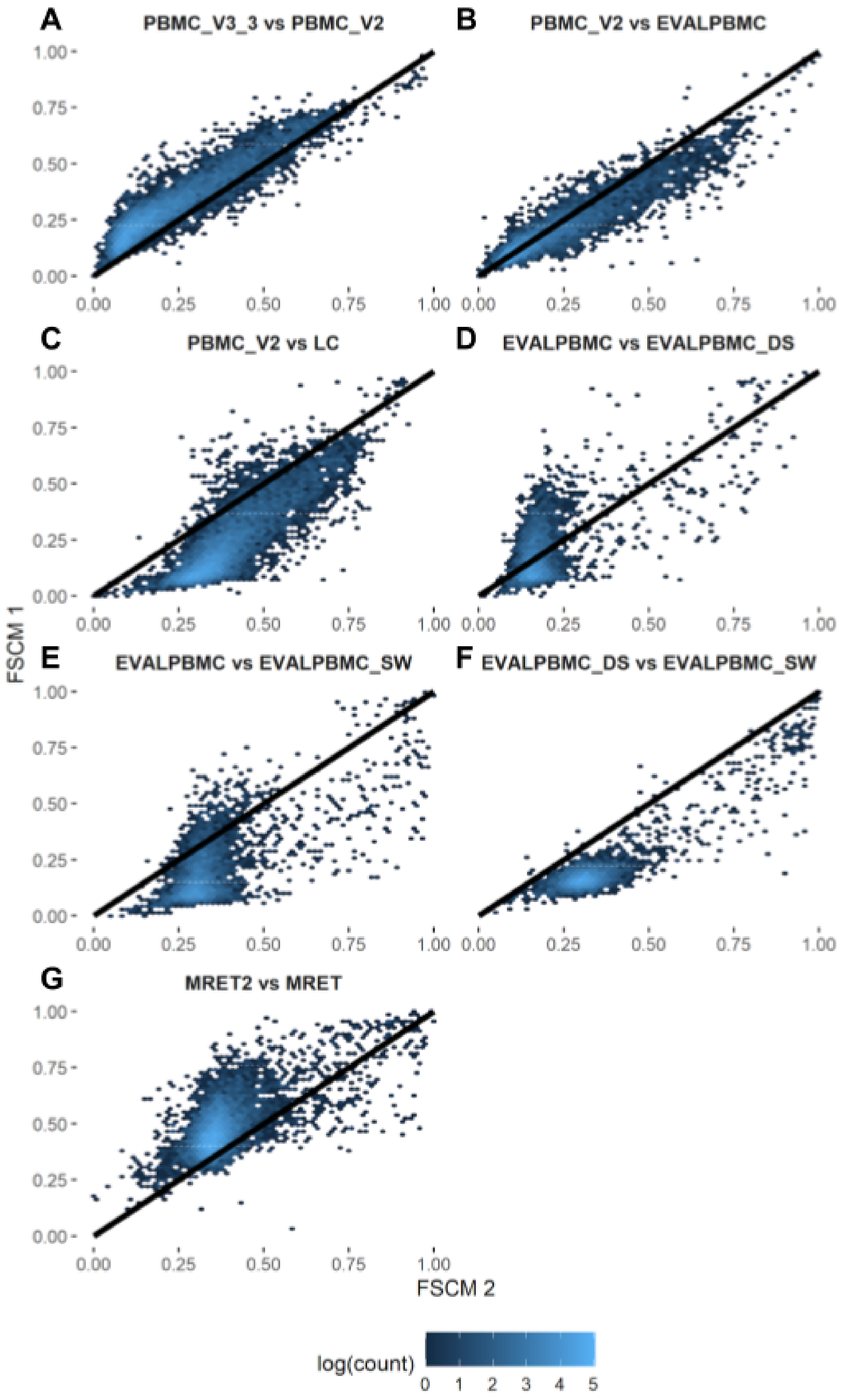

A
PBMC_V3_3 vs PBMC_V2
B
PBMC_V2 vs EVALPBMC
C
PBMC_V2 vs LC
D
EVALPBMC vs EVALPBMC_DS
E
EVALPBMC vs EVALPBMC_SW
F
EVALPBMC_DS vs EVALPBMC_SW
G
MRET2 vs MRET
0.00
0.25
0.50
0.75
1.00
FSCM 1
FSCM 2
log(count)
0
1
2
3
4
5

***Fig S5: Gene-wise comparison of FSCM between pairs of datasets.*** *Each point represents a gene; only genes with at least 200 UMIs in both datasets are included. FSCM 1 (y axis) refers to the first dataset in the title and FSCM 2 (x axis) to the second. A. Both datasets from 10x Chromium, v3 chemistry, human PBMC (identical to Fig. 2B in the main text). B. 10x Chromium, v2 vs v3 chemistry, human PBMC. C. Both datasets from 10x Chromium, v2 chemistry, human PBMC, but from different labs. D. Both datasets from 10x Chromium, v2 chemistry, but from different labs and different tissue - human PBMC vs human lung tumor. E. 10x Chromium, v2 chemistry, vs Drop-Seq, both human PBMC, generated from the same sample. F. 10x Chromium, v2 chemistry, vs Seq-Well, both human PBMC, generated from the same sample. G. Drop-Seq vs Seq-Well, both human PBMC, generated from the same sample. H. 10x Chromium, v2 chemistry, vs Drop-Seq, both mouse retina but generated by different research groups.*
*The code to reproduce this figure is here: code*

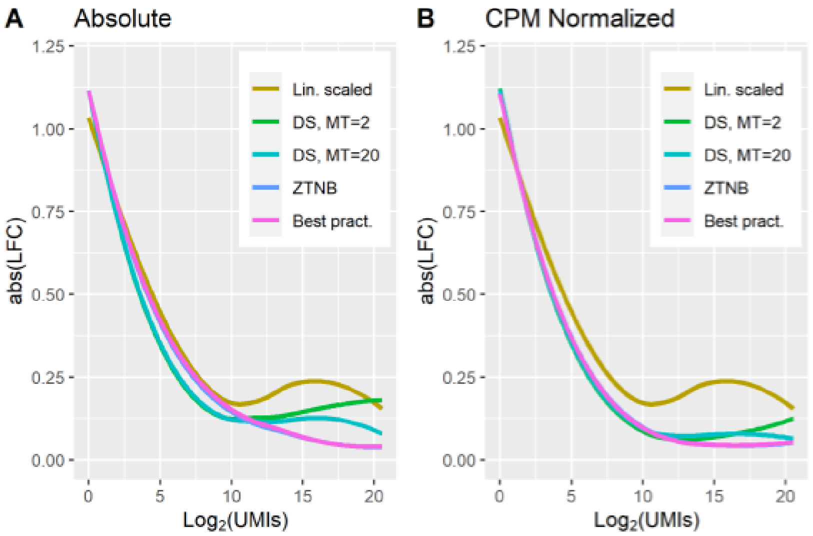


| Method | CCC |
|---|---|
| Lin. Scaled | 0.9810 |
| DS, MT=2 | 0.9905 |
| DS, MT=20 | 0.9901 |
| ZTNB | 0.9899 |
| Best pract. | 0.9899 |

***Fig S6:*** *Prediction errors for different methods. Each curve represents a loess fit of abs($Log_2$ Fold Change) over all genes and all datasets, where the predicted value is compared to ground truth. A. Direct comparison of predicted molecules per gene. B. Comparison of predicted molecules per gene after CPM normalization. The normalization removes systematic errors in prediction across genes, for example if a method underestimates all predictions. The CCC values presented are calculated on log-transformed CPM-normalized data. The code to reproduce this figure is here:* [*code*]

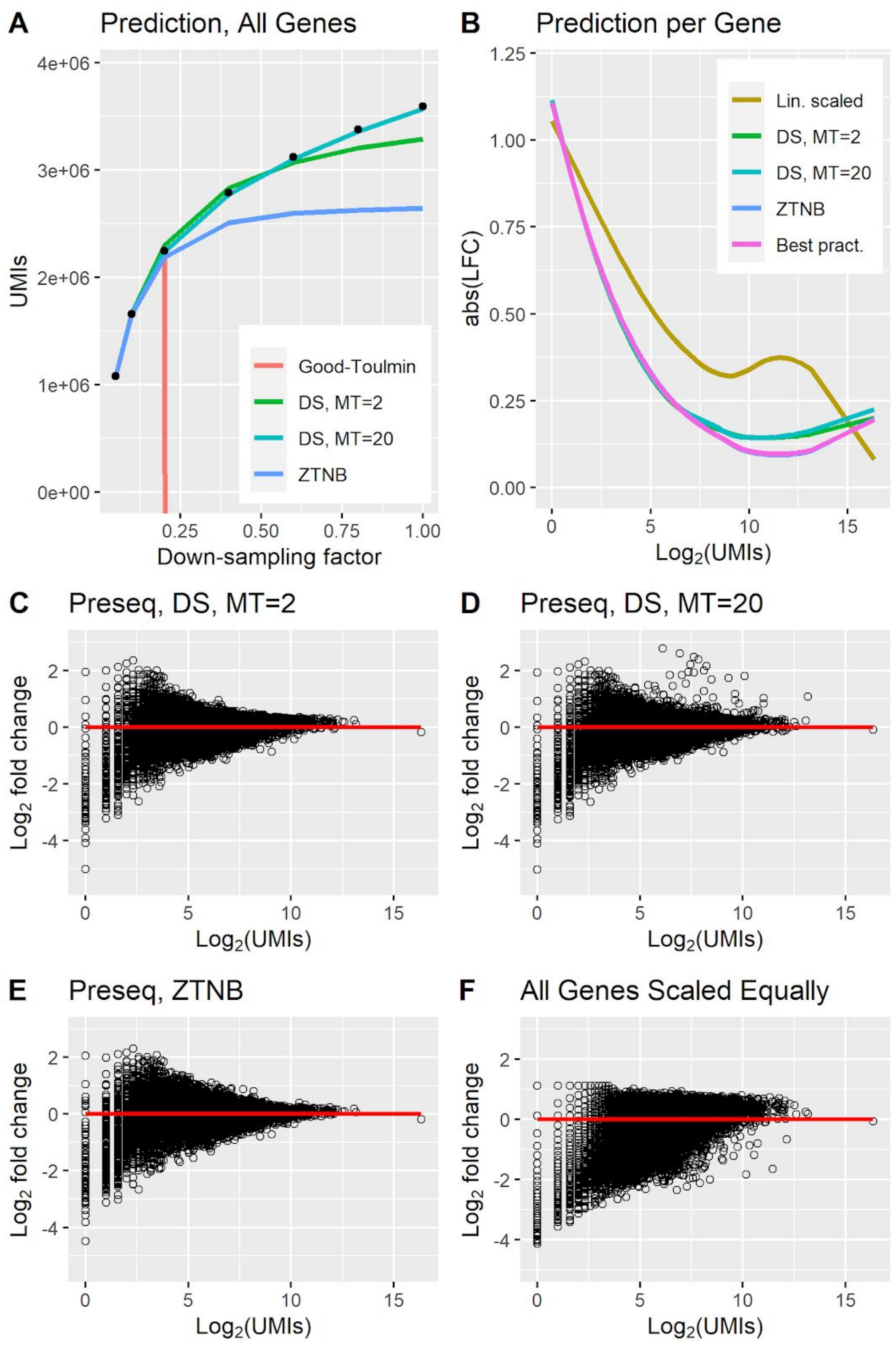


***Fig S7: Correction evaluation for the EVAL dataset.*** *The data was downsampled to 1/20 for A and to 1/10 for B-F, and the corrected expression using different prediction methods was compared to ground truth (predicting to 10 times the number of counts for B-F). A. Prediction of all UMIs in the dataset, collected into a single pool. The data was corrected from 1/20 of the reads. Ground truth is represented by black dots. B. Correction errors for different prediction methods on CPM-normalized data. The figure shows a loess fit of abs(LFC) over all genes. C-F. Scatter plots showing the correction error for each gene as the* $Log_2$ *fold change between corrected expression and ground truth (CPM normalized). The x axis corresponds to* $Log_2$ *of the number of UMIs for the gene in the downsampled data. The code to reproduce this figure is here: code*

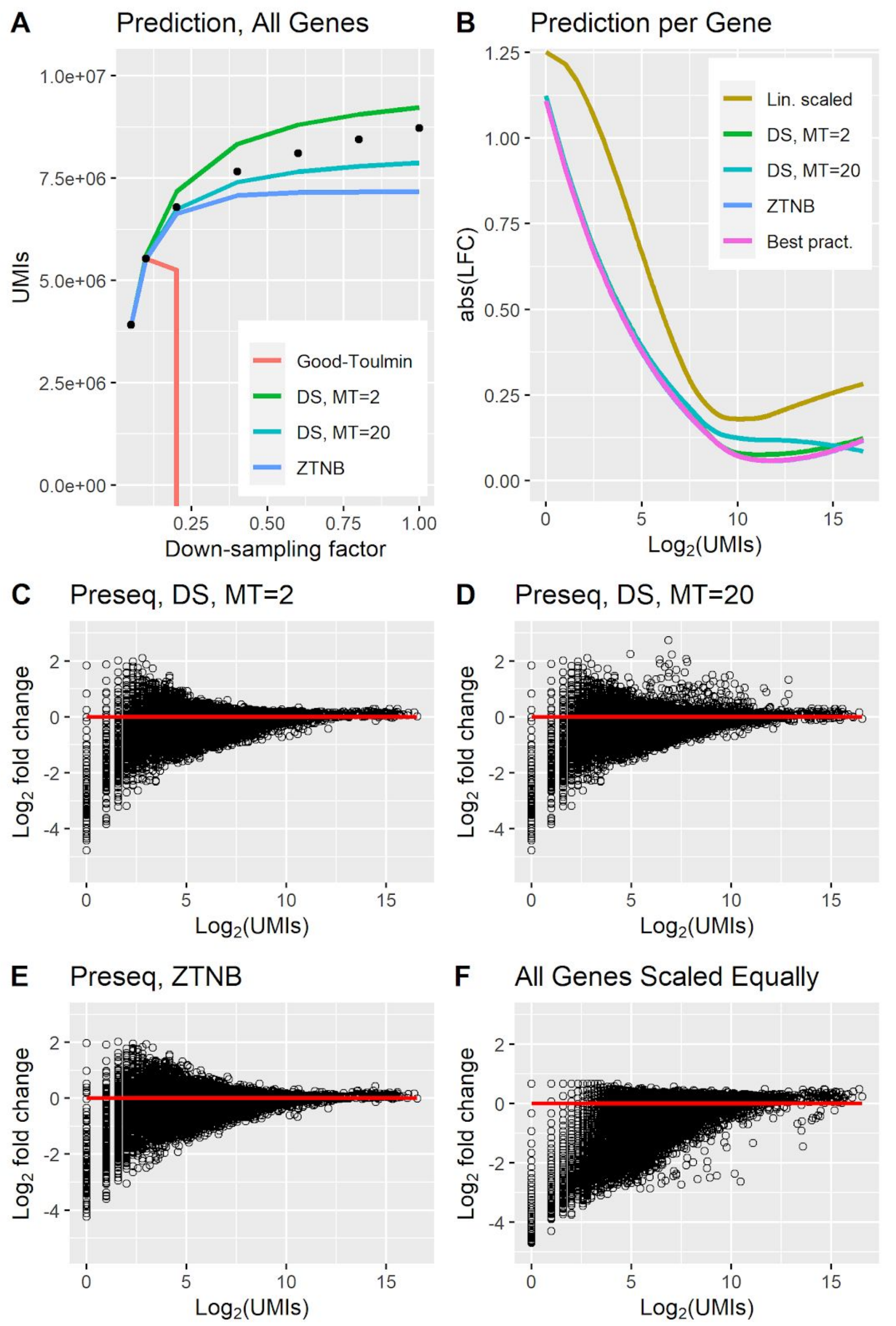


***Fig S8: Correction evaluation for the EVALPBMC dataset.*** *The data was downsampled to 1/20 for A and to 1/10 for B-F, and the corrected expression using different prediction methods was compared to ground truth (predicting to 10 times the number of counts for B-F). A. Prediction of all UMIs in the dataset, collected into a single pool. The data was corrected from 1/20 of the reads. Ground truth is represented by black dots. B. Correction errors for different prediction methods on CPM-normalized data. The figure shows a loess fit of abs(LFC) over all genes. C-F. Scatter plots showing the correction error for each gene as the* $Log_2$ *fold change between corrected expression and ground truth (CPM normalized). The x axis corresponds to* $Log_2$ *of the number of UMIs for the gene in the downsampled data. The code to reproduce this figure is here:* [code]

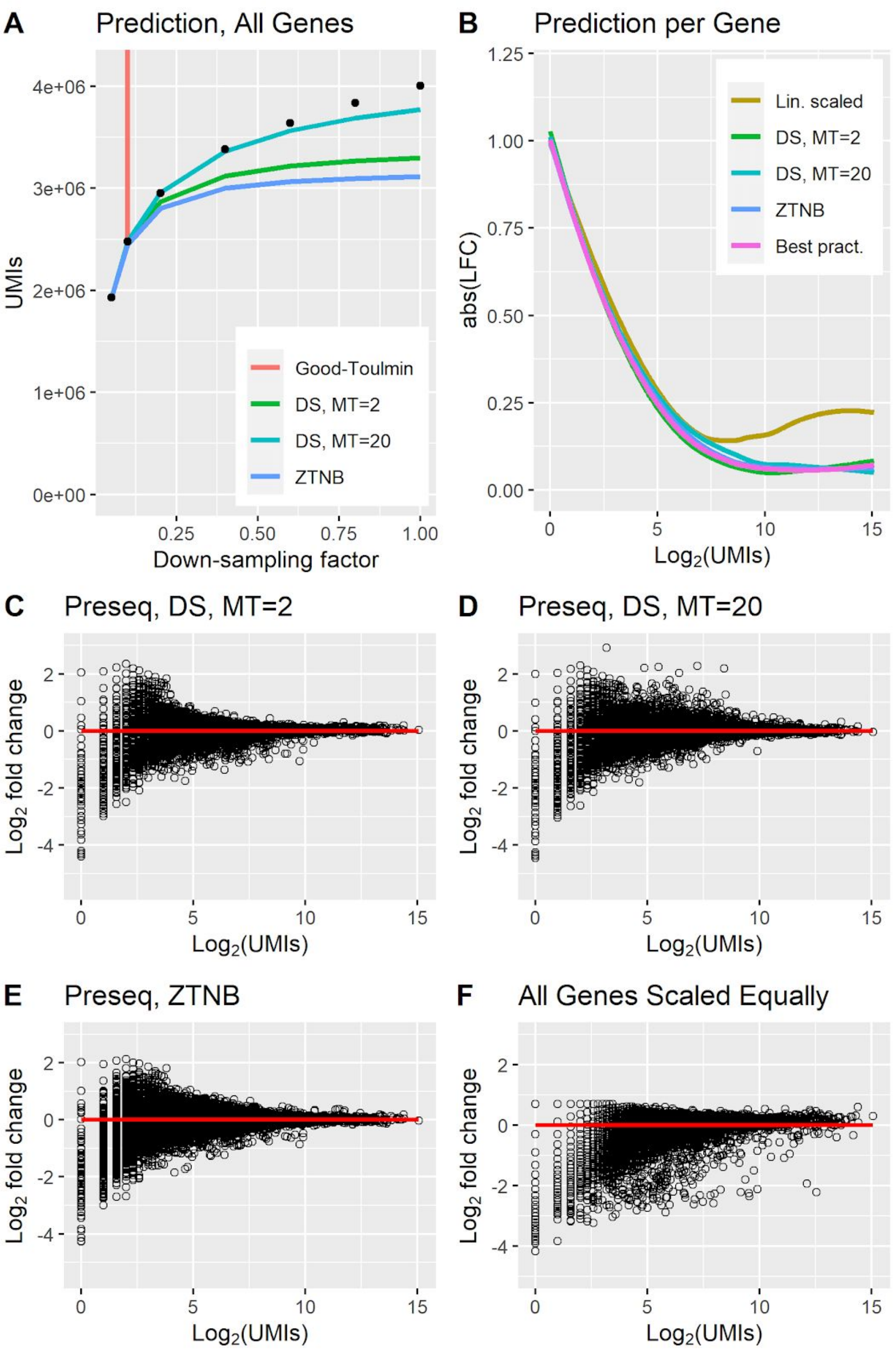


***Fig S9: Correction evaluation for the EVALPBMC_DS dataset.*** *The data was downsampled to 1/20 for A and to 1/10 for B-F, and the corrected expression using different prediction methods was compared to ground truth (predicting to 10 times the number of counts for B-F). A. Prediction of all UMIs in the dataset, collected into a single pool. The data was corrected from 1/20 of the reads. Ground truth is represented by black dots. B. Correction errors for different prediction methods on CPM-normalized data. The figure shows a loess fit of abs(LFC) over all genes. C-F. Scatter plots showing the correction error for each gene as the* $Log_2$ *fold change between corrected expression and ground truth (CPM normalized). The x axis corresponds to* $Log_2$ *of the number of UMIs for the gene in the downsampled data. The code to reproduce this figure is here:* [code]

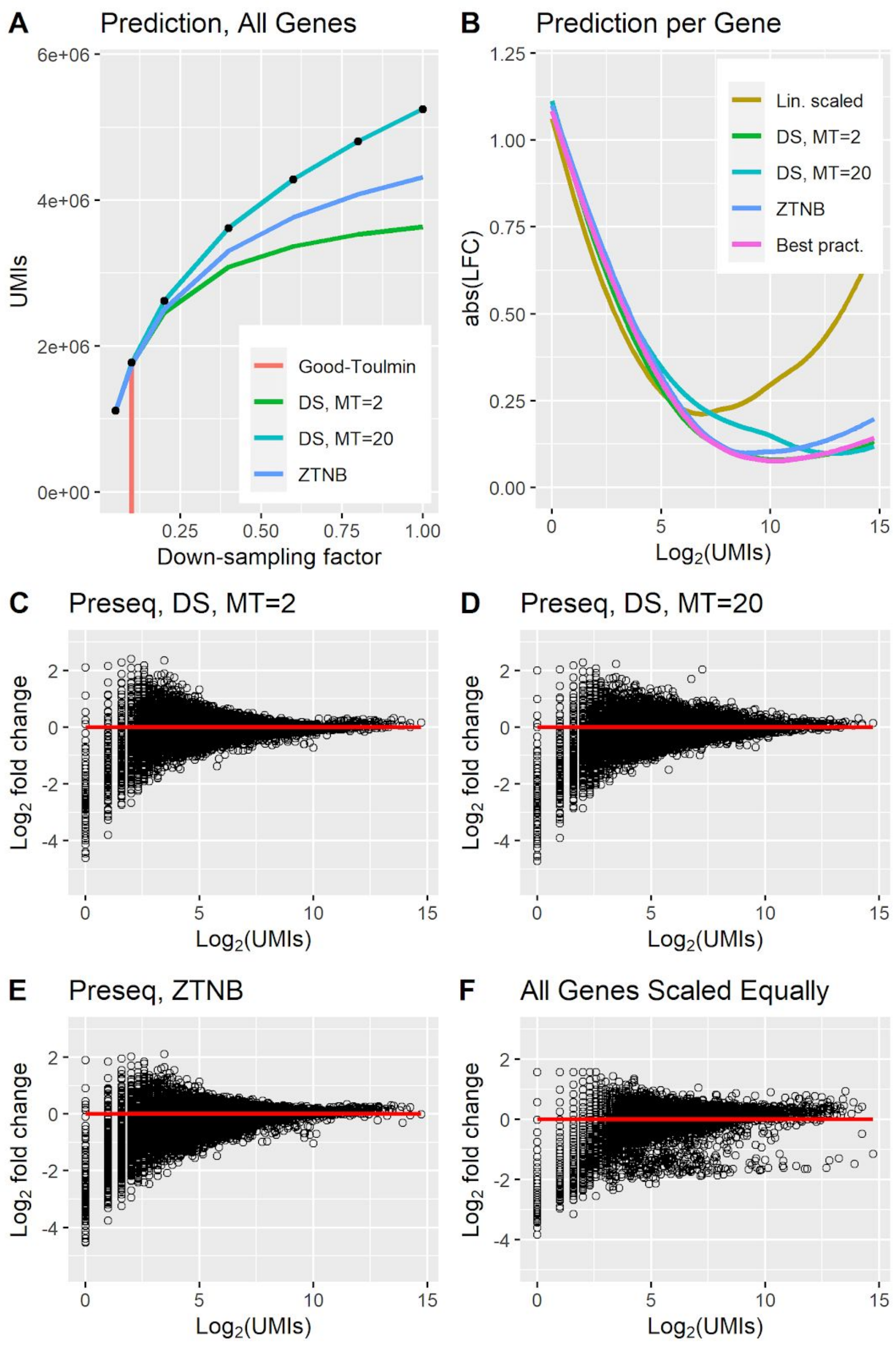


***Fig S10: Correction evaluation for the EVALPBMC_SW dataset.*** *The data was downsampled to 1/20 for A and to 1/10 for B-F, and the corrected expression using different prediction methods was compared to ground truth (predicting to 10 times the number of counts for B-F). A. Prediction of all UMIs in the dataset, collected into a single pool. The data was corrected from 1/20 of the reads. Ground truth is represented by black dots. B. Correction errors for different prediction methods on CPM-normalized data. The figure shows a loess fit of abs(LFC) over all genes. C-F. Scatter plots showing the correction error for each gene as the $Log_2$ fold change between corrected expression and ground truth (CPM normalized). The x axis corresponds to $Log_2$ of the number of UMIs for the gene in the downsampled data. The code to reproduce this figure is here:* [*code*]

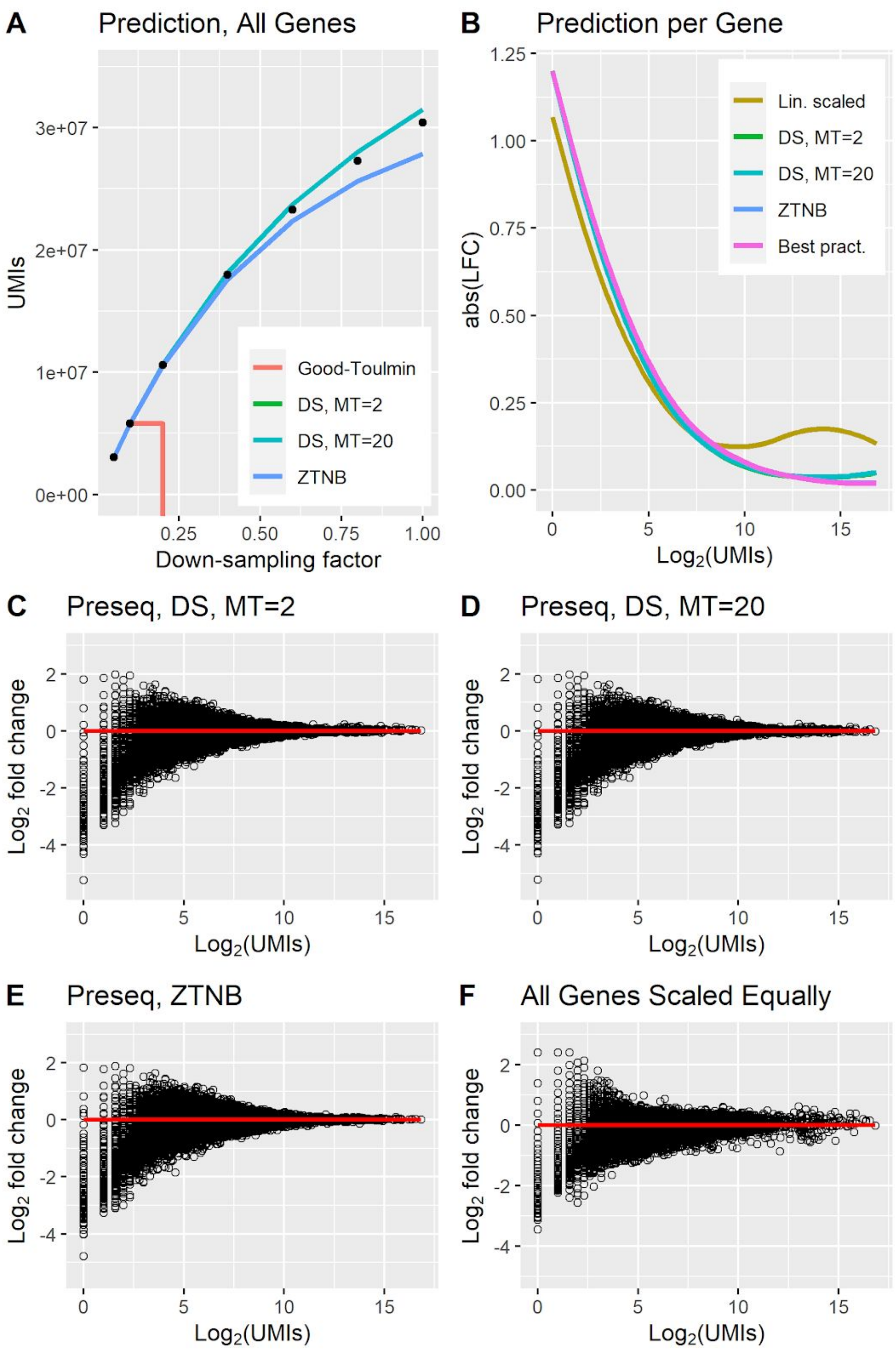


***Fig S11: Correction evaluation for the PBMC_V3 dataset.*** *The data was downsampled to 1/20 for A and to 1/10 for B-F, and the corrected expression using different prediction methods was compared to ground truth (predicting to 10 times the number of counts for B-F). A. Prediction of all UMIs in the dataset, collected into a single pool. The data was corrected from 1/20 of the reads. Ground truth is represented by black dots. B. Correction errors for different prediction methods on CPM-normalized data. The figure shows a loess fit of abs(LFC) over all genes. C-F. Scatter plots showing the correction error for each gene as the $Log_2$ fold change between corrected expression and ground truth (CPM normalized). The x axis corresponds to $Log_2$ of the number of UMIs for the gene in the downsampled data. The code to reproduce this figure is here: [code]*

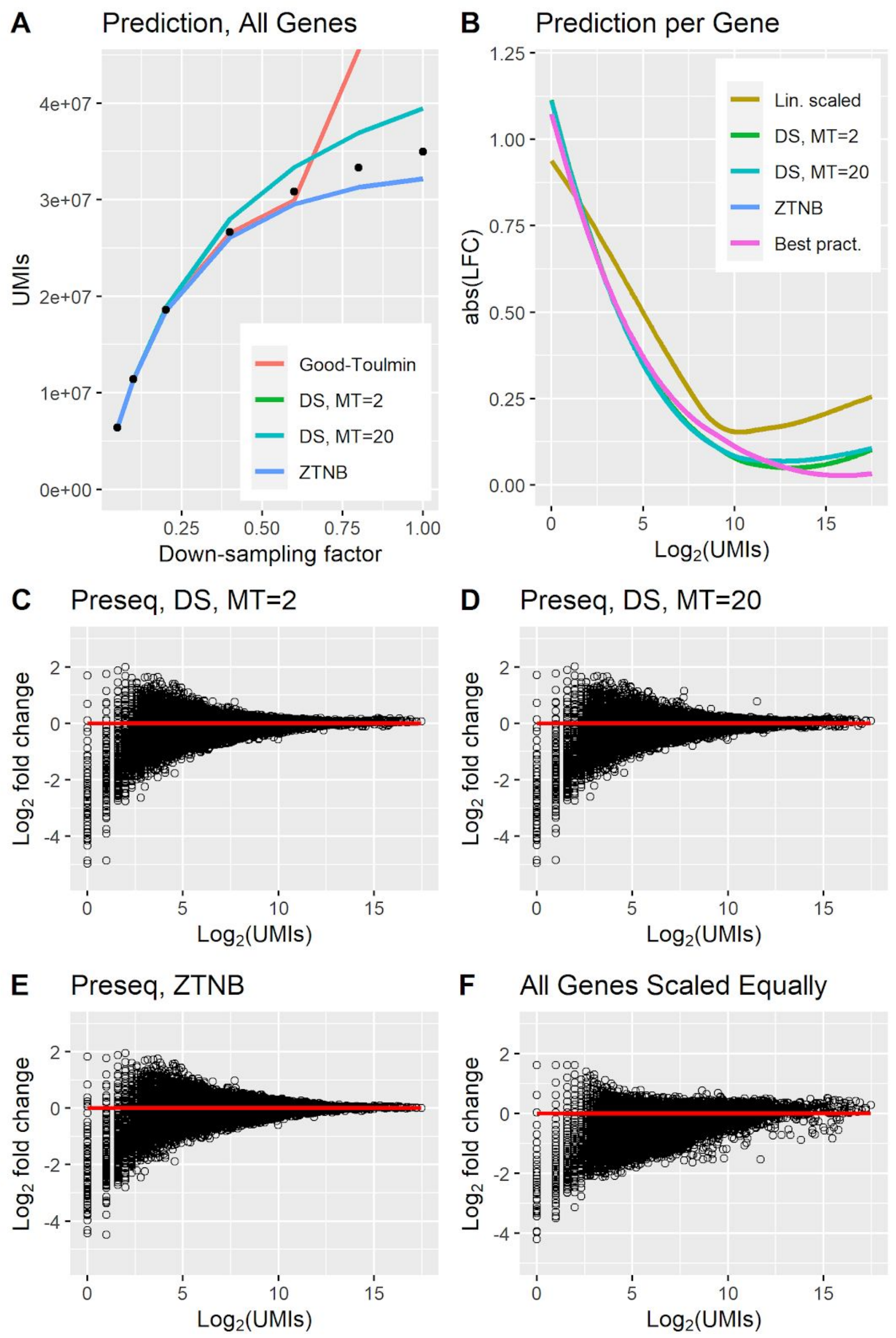


***Fig S12: Correction evaluation for the PBMC_V3_2 dataset.*** *The data was downsampled to 1/20 for A and to 1/10 for B-F, and the corrected expression using different prediction methods was compared to ground truth (predicting to 10 times the number of counts for B-F). A. Prediction of all UMIs in the dataset, collected into a single pool. The data was corrected from 1/20 of the reads. Ground truth is represented by black dots. B. Correction errors for different prediction methods on CPM-normalized data. The figure shows a loess fit of abs(LFC) over all genes. C-F. Scatter plots showing the correction error for each gene as the* $Log_2$ *fold change between corrected expression and ground truth (CPM normalized). The x axis corresponds to* $Log_2$ *of the number of UMIs for the gene in the downsampled data. The code to reproduce this figure is here: code*

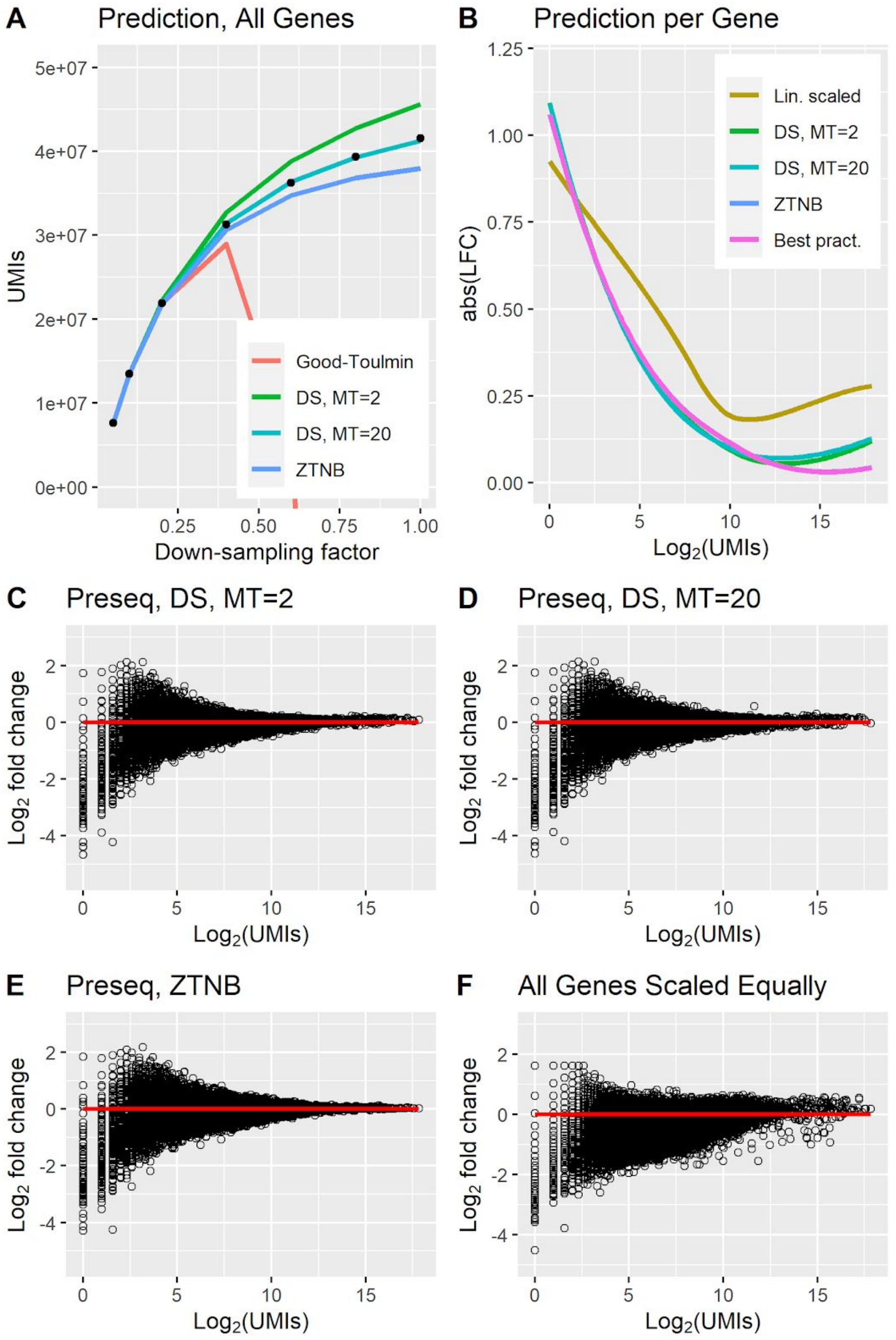


***Fig S13: Correction evaluation for the PBMC_V3_3 dataset.*** *The data was downsampled to 1/20 for A and to 1/10 for B-F, and the corrected expression using different prediction methods was compared to ground truth (predicting to 10 times the number of counts for B-F). A. Prediction of all UMIs in the dataset, collected into a single pool. The data was corrected from 1/20 of the reads. Ground truth is represented by black dots. B. Correction errors for different prediction methods on CPM-normalized data. The figure shows a loess fit of abs(LFC) over all genes. C-F. Scatter plots showing the correction error for each gene as the $Log_2$ fold change between corrected expression and ground truth (CPM normalized). The x axis corresponds to $Log_2$ of the number of UMIs for the gene in the downsampled data. The code to reproduce this figure is here: code*

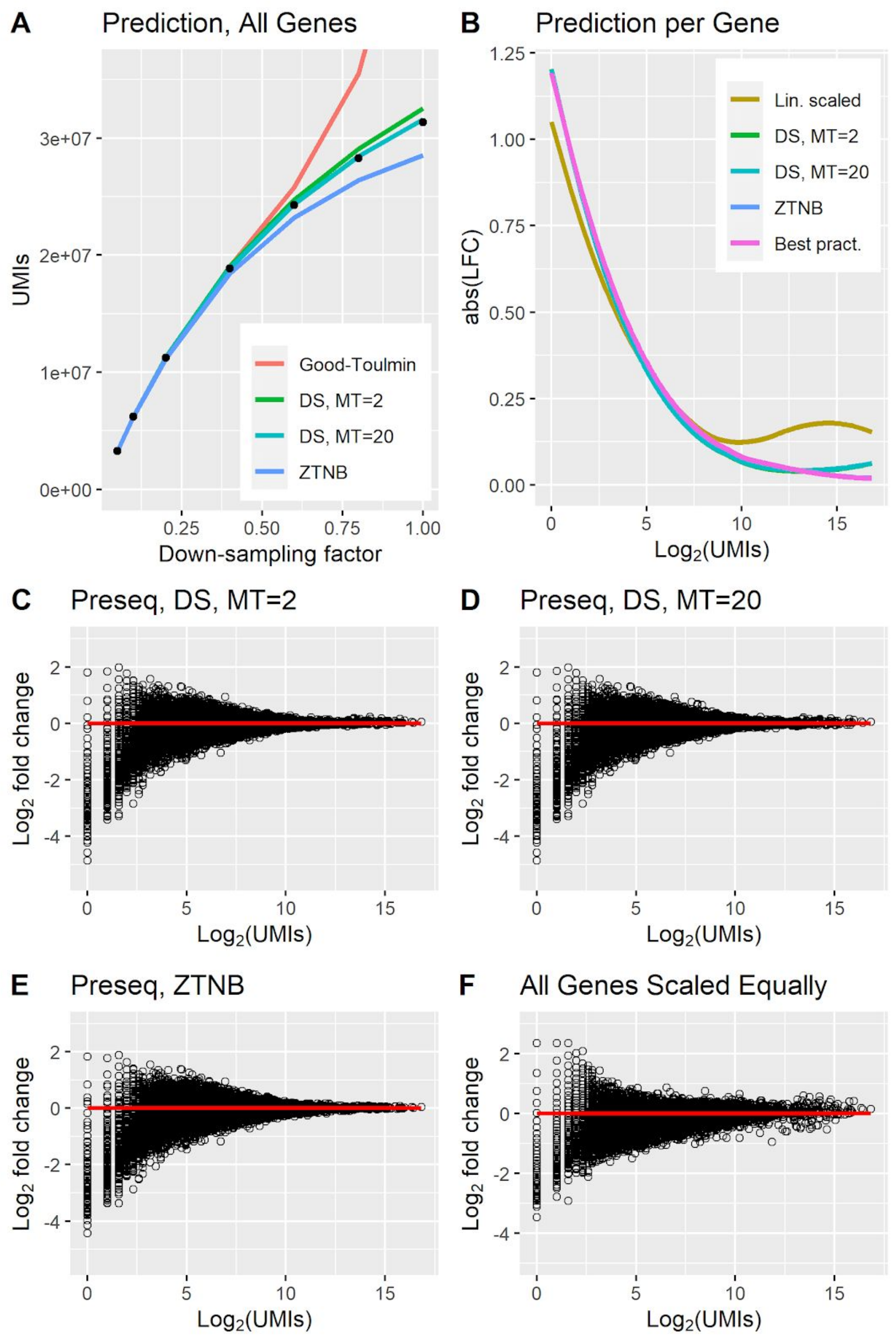


***Fig S14: Correction evaluation for the PBMC_NG dataset.*** *The data was downsampled to 1/20 for A and to 1/10 for B-F, and the corrected expression using different prediction methods was compared to ground truth (predicting to 10 times the number of counts for B-F). A. Prediction of all UMIs in the dataset, collected into a single pool. The data was corrected from 1/20 of the reads. Ground truth is represented by black dots. B. Correction errors for different prediction methods on CPM-normalized data. The figure shows a loess fit of abs(LFC) over all genes. C-F. Scatter plots showing the correction error for each gene as the $Log_2$ fold change between corrected expression and ground truth (CPM normalized). The x axis corresponds to $Log_2$ of the number of UMIs for the gene in the downsampled data. The code to reproduce this figure is here: [code]*

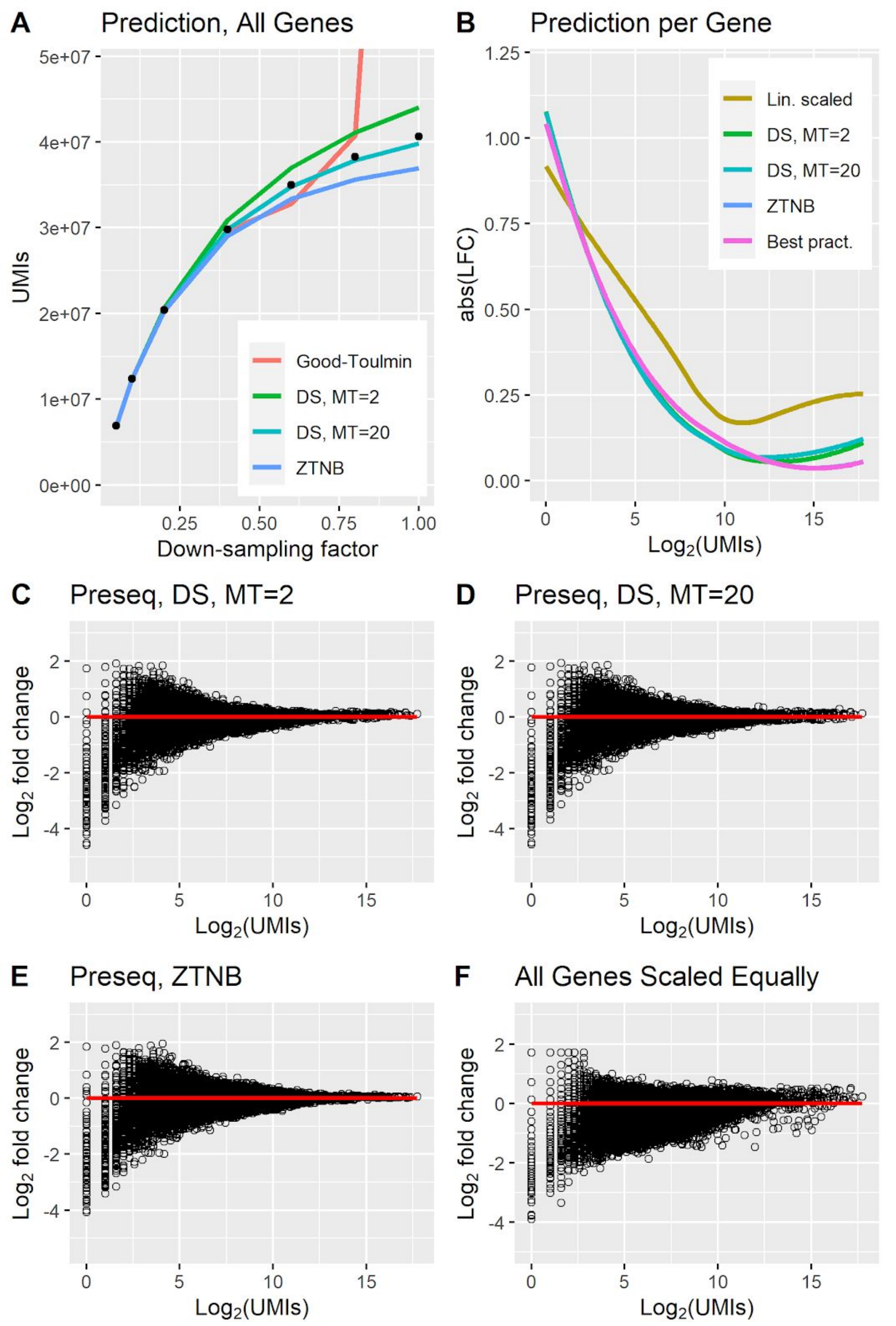


***Fig S15: Correction evaluation for the PBMC_NG_2 dataset.*** *The data was downsampled to 1/20 for A and to 1/10 for B-F, and the corrected expression using different prediction methods was compared to ground truth (predicting to 10 times the number of counts for B-F). A. Prediction of all UMIs in the dataset, collected into a single pool. The data was corrected from 1/20 of the reads. Ground truth is represented by black dots. B. Correction errors for different prediction methods on CPM-normalized data. The figure shows a loess fit of abs(LFC) over all genes. C-F. Scatter plots showing the correction error for each gene as the* $Log_2$ *fold change between corrected expression and ground truth (CPM normalized). The x axis corresponds to* $Log_2$ *of the number of UMIs for the gene in the downsampled data. The code to reproduce this figure is here: [code]*

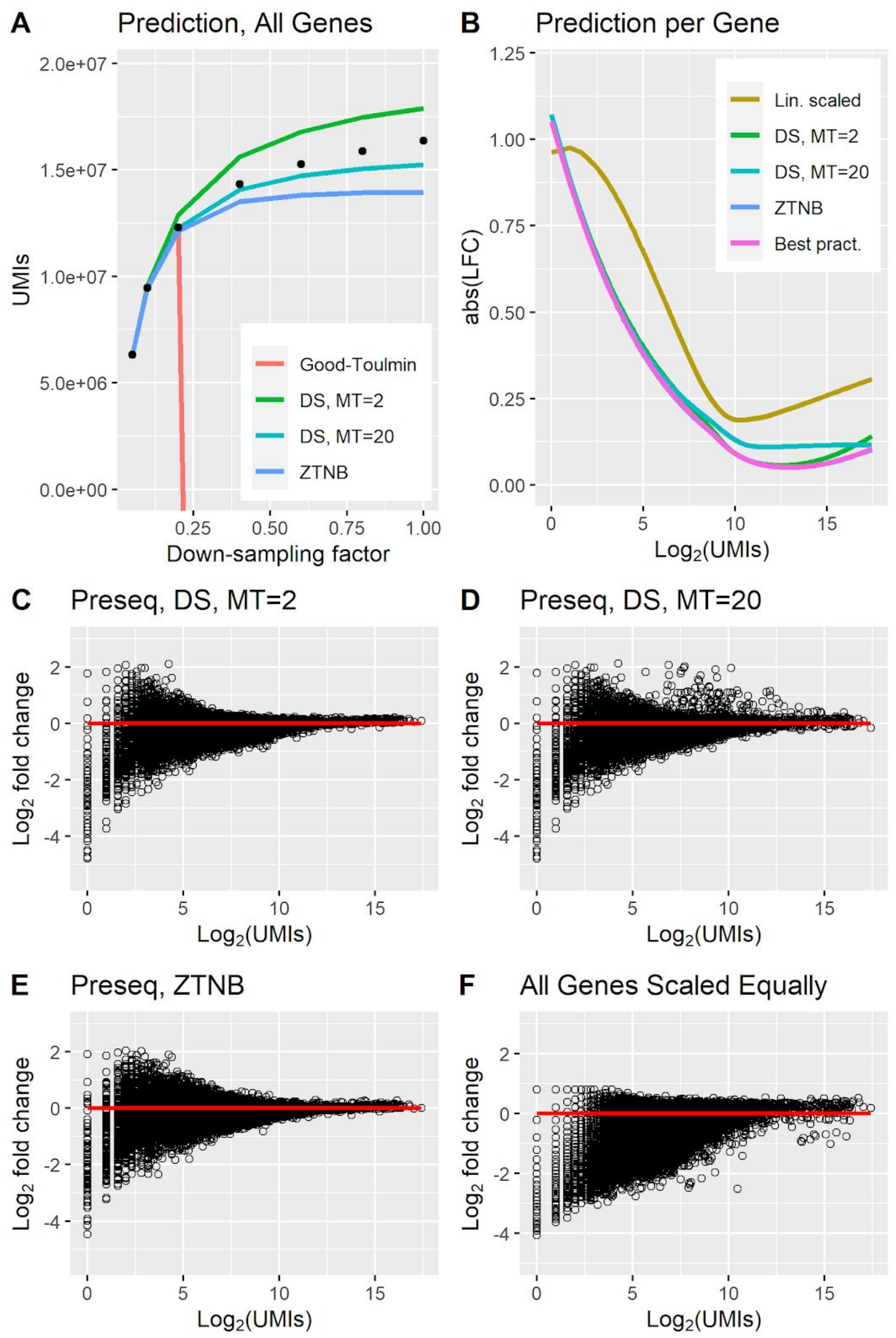


***Fig S16: Correction evaluation for the PBMC_V2 dataset.*** *The data was downsampled to 1/20 for A and to 1/10 for B-F, and the corrected expression using different prediction methods was compared to ground truth (predicting to 10 times the number of counts for B-F). A. Prediction of all UMIs in the dataset, collected into a single pool. The data was corrected from 1/20 of the reads. Ground truth is represented by black dots. B. Correction errors for different prediction methods on CPM-normalized data. The figure shows a loess fit of abs(LFC) over all genes. C-F. Scatter plots showing the correction error for each gene as the $Log_2$ fold change between corrected expression and ground truth (CPM normalized). The x axis corresponds to $Log_2$ of the number of UMIs for the gene in the downsampled data. The code to reproduce this figure is here: [code]*

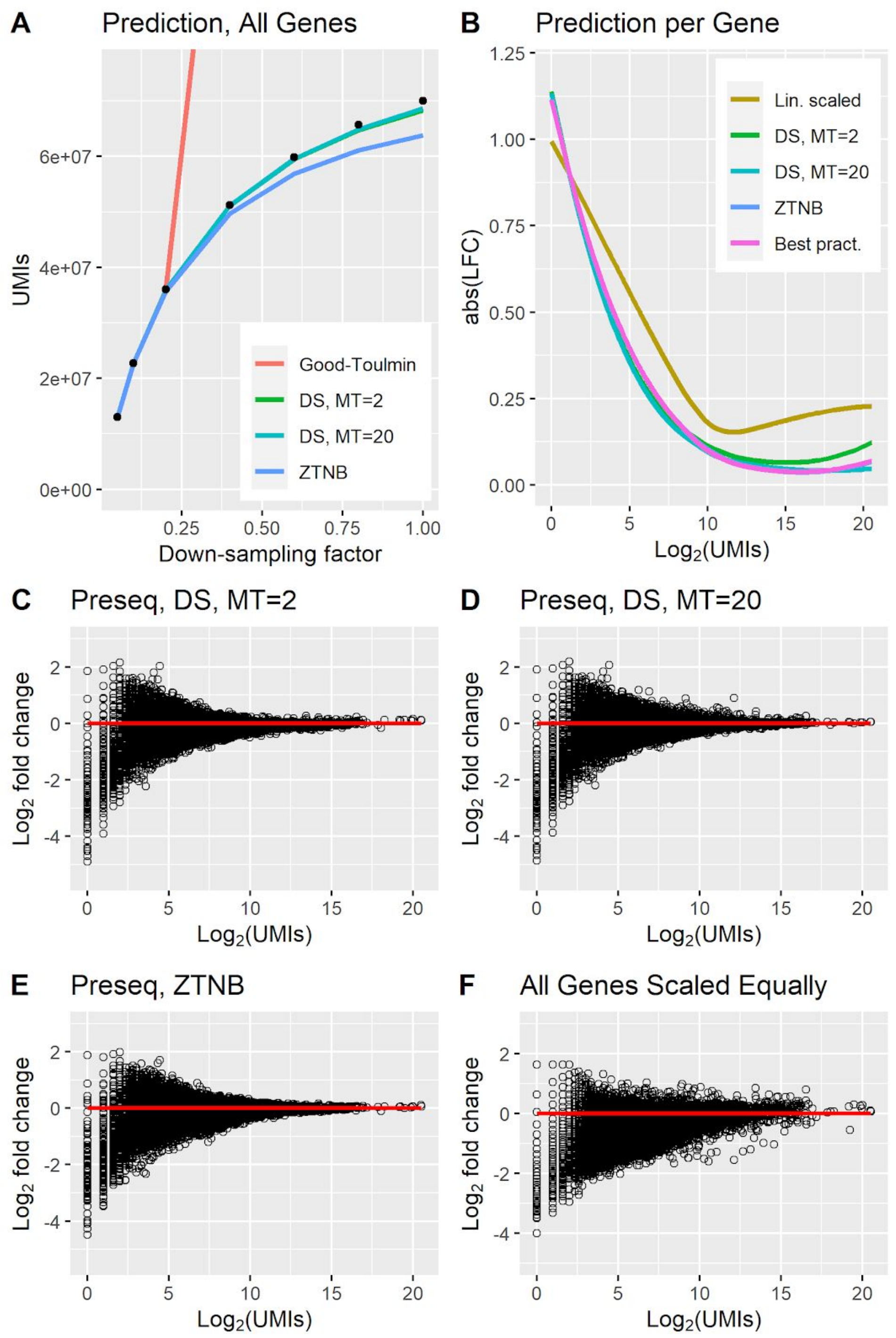


***Fig S17: Correction evaluation for the LC dataset.*** *The data was downsampled to 1/20 for A and to 1/10 for B-F, and the corrected expression using different prediction methods was compared to ground truth (predicting to 10 times the number of counts for B-F). A. Prediction of all UMIs in the dataset, collected into a single pool. The data was corrected from 1/20 of the reads. Ground truth is represented by black dots. B. Correction errors for different prediction methods on CPM-normalized data. The figure shows a loess fit of abs(LFC) over all genes. C-F. Scatter plots showing the correction error for each gene as the $Log_2$ fold change between corrected expression and ground truth (CPM normalized). The x axis corresponds to $Log_2$ of the number of UMIs for the gene in the downsampled data.*
*The code to reproduce this figure is here: [code]*

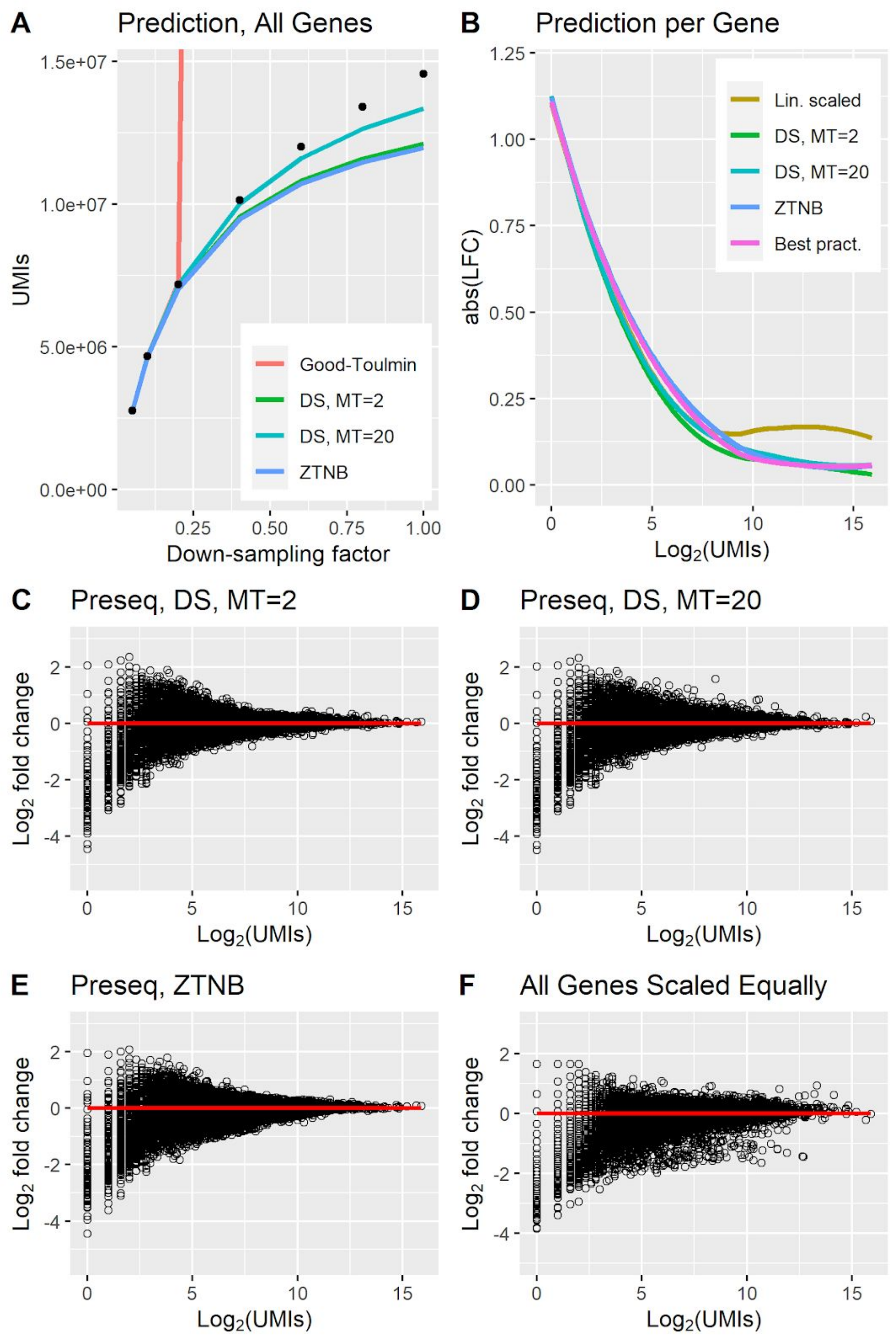


***Fig S18: Correction evaluation for the MRET dataset.*** *The data was downsampled to 1/20 for A and to 1/10 for B-F, and the corrected expression using different prediction methods was compared to ground truth (predicting to 10 times the number of counts for B-F). A. Prediction of all UMIs in the dataset, collected into a single pool. The data was corrected from 1/20 of the reads. Ground truth is represented by black dots. B. Correction errors for different prediction methods on CPM-normalized data. The figure shows a loess fit of abs(LFC) over all genes. C-F. Scatter plots showing the correction error for each gene as the $Log_2$ fold change between corrected expression and ground truth (CPM normalized). The x axis corresponds to $Log_2$ of the number of UMIs for the gene in the downsampled data. The code to reproduce this figure is here:* [*code*]

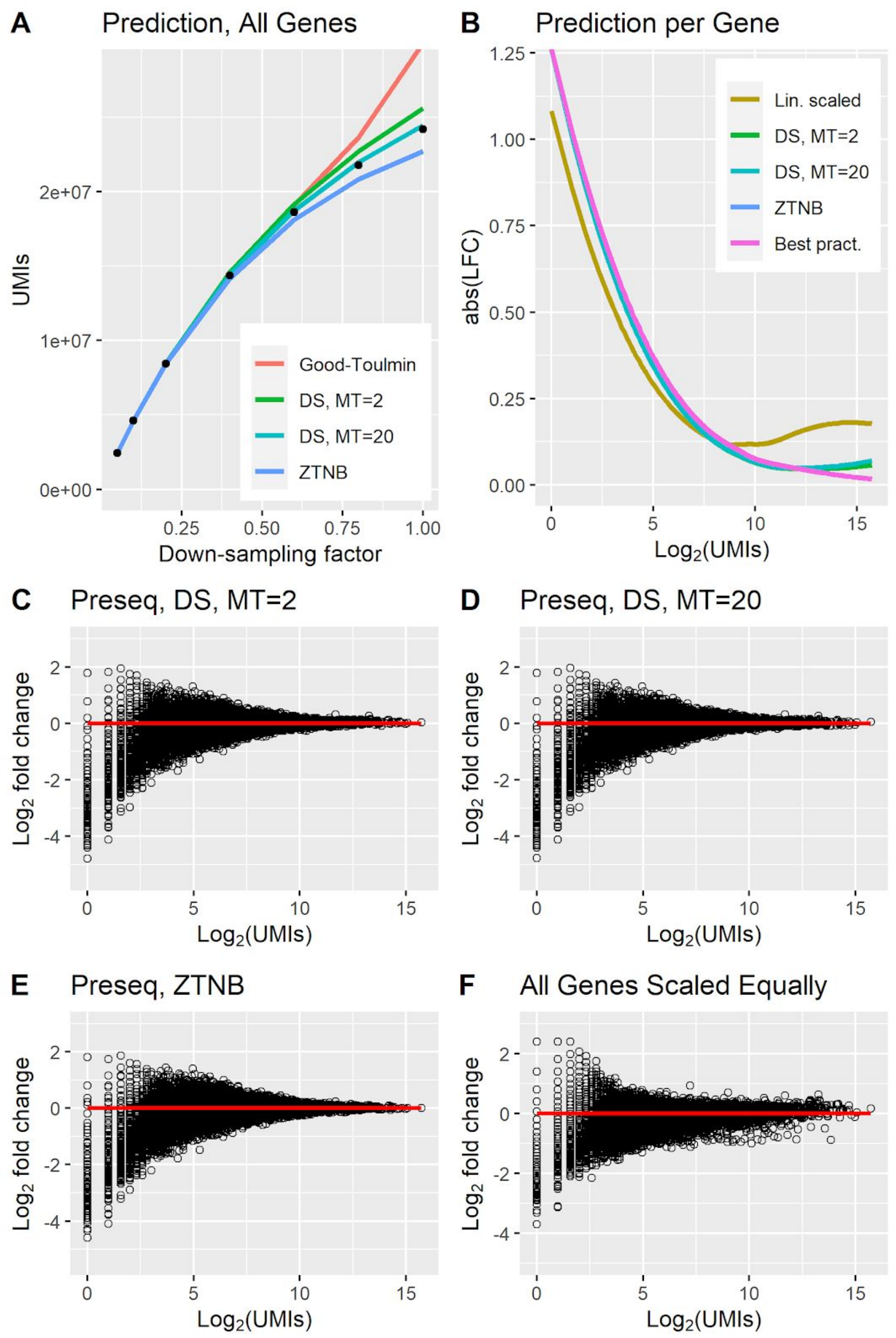


***Fig S19: Correction evaluation for the MRET2 dataset.*** *The data was downsampled to 1/20 for A and to 1/10 for B-F, and the corrected expression using different prediction methods was compared to ground truth (predicting to 10 times the number of counts for B-F). A. Prediction of all UMIs in the dataset, collected into a single pool. The data was corrected from 1/20 of the reads. Ground truth is represented by black dots. B. Correction errors for different prediction methods on CPM-normalized data. The figure shows a loess fit of abs(LFC) over all genes. C-F. Scatter plots showing the correction error for each gene as the* $Log_2$ *fold change between corrected expression and ground truth (CPM normalized). The x axis corresponds to* $Log_2$ *of the number of UMIs for the gene in the downsampled data. The code to reproduce this figure is here: code*

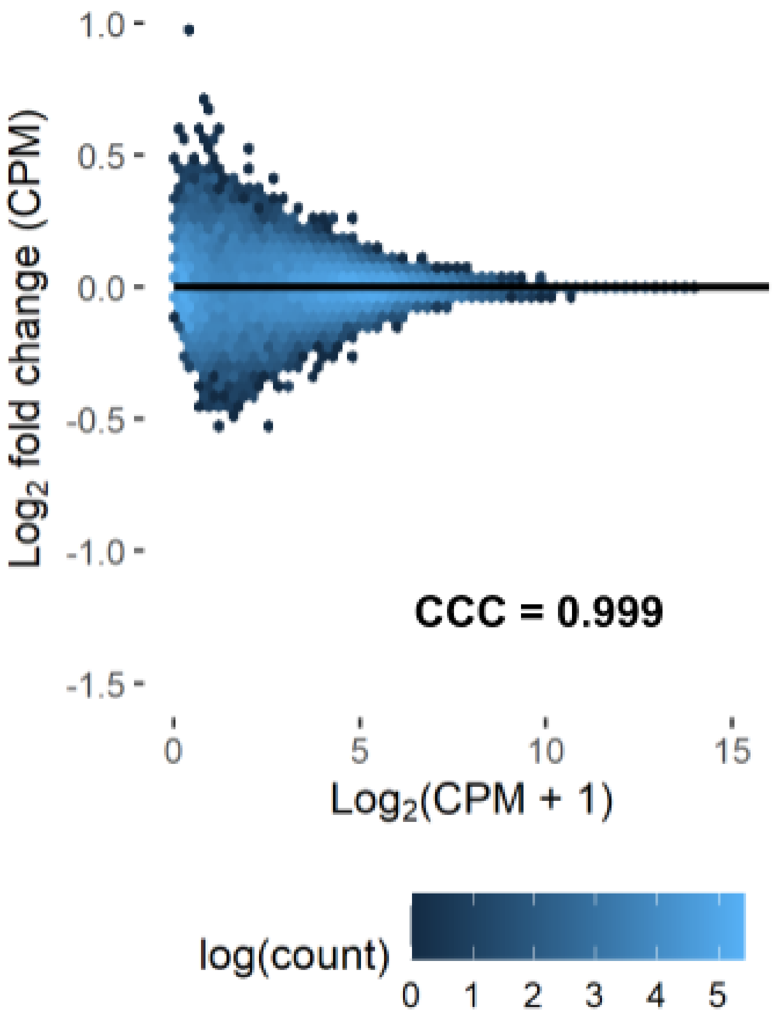


***Fig S20: Sampling noise from downsampling.*** *Downsampled data compared to the mean expression of downsampling 20 times. No amplification bias is present here, since the gene expressions compared have the same number of reads per cell. This provides a bound on the accuracy possible with correction of unseen molecules in downsampled data.The code to reproduce this figure is here: code*

### Supplementary Tables

| Id | Description | Technology | Reference |
|---|---|---|---|
| EVAL | Mouse brain | 10x Chromium, v2 | Ding et al(1), GSE132044, Cortex 1 |
| EVALPBMC | Human PBMC | 10x Chromium, v2 | Ding et al(1), GSE132044, PBMC 1 |
| EVALPBMC_DS | Human PBMC, same sample as EVALPBMC | Drop-Seq | Ding et al(1), GSE132044, PBMC 1 |
| EVALPBMC_SW | Human PBMC, same sample as EVALPBMC | Seq-Well | Ding et al(1), GSE132044, PBMC 1 |
| PBMC_V3 | Human PBMC | 10x Chromium, v3 | Available at 10x Genomics' home page(2). Protein data not used. |
| PBMC_V3_2 | Human PBMC | 10x Chromium, v3 | Available at 10x Genomics' home page(3). Protein data not used. |
| PBMC_V3_3 | Human PBMC | 10x Chromium, v3 | Available at 10x Genomics' home page(4). |
| PBMC_NG | Human PBMC | 10x Chromium, NetxGEM | Available at 10x Genomics' home page(5). Protein data not used. |
| PBMC_NG_2 | Human PBMC, same sample as PBMC_V3_3. | 10x Chromium, NetxGEM | Available at 10x Genomics' home page(6). |
| PBMC_V2 | Human PBMC | 10x Chromium, v2 | Available at 10x Genomics' home page(7). |
| LC | Lung tumor | 10x Chromium, v2 | Lambrechts et al(8). The data is available in ArrayExpress under the ascension E-MTAB-6149. Data for Patient 3, combined edge, middle and core of the tumor, was used. |
| MRET | Mouse retina | Drop-Seq | Macosco et al(9), GSE63473, P14 mouse retina 7. |
| MRET2 | Mouse retina | 10x Chromium, v2 | Clark et al(10), GSE117614, sample p0. |

*Table S1. List of datasets used in this study.*

| Id | Cells (k) | Tot Cnts (M) | Tot UMIs (M) | Avg CU | Avg UMIs per Cell (k) | Avg Cnts per Cell (k) | Tot FSCM |
|---|---|---|---|---|---|---|---|
| EVAL | 1.6 | 30 | 3.6 | 8.30 | 2.3 | 19.2 | 0.277 |
| EVAL_PBMC | 5.2 | 112 | 8.7 | 12.9 | 1.7 | 21.6 | 0.147 |
| EVAL_PBMC_DS | 6.9 | 87 | 4.0 | 21.8 | 0.59 | 12.7 | 0.199 |
| EVAL_PBMC_SW | 7.2 | 32 | 5.3 | 6.05 | 0.73 | 4.44 | 0.391 |
| PBMC_V3 | 5.4 | 64 | 30 | 2.11 | 5.6 | 11.8 | 0.453 |
| PBMC_V3_2 | 8.2 | 144 | 35 | 4.12 | 4.3 | 17.5 | 0.199 |
| PBMC_V3_3 | 5.2 | 173 | 42 | 4.16 | 7.9 | 33.0 | 0.225 |
| PBMC_NG | 5.6 | 69 | 31 | 2.19 | 5.6 | 12.2 | 0.435 |
| PBMC_NG_2 | 5.4 | 155 | 41 | 3.81 | 7.6 | 28.9 | 0.248 |
| PBMC_V2 | 4.9 | 180 | 16 | 11.0 | 3.4 | 36.9 | 0.131 |
| LC | 21.1 | 305 | 70 | 4.36 | 3.3 | 14.4 | 0.267 |
| MRET | 20.2 | 68 | 15 | 4.67 | 0.72 | 3.4 | 0.367 |
| MRET2 | 12.5 | 51 | 24 | 2.09 | 1.93 | 4.1 | 0.440 |

*Table S2. Statistics for the datasets used in the study. FSCM is the fraction of single-copy molecules.*

### Supplementary Note

We evaluated 2 different algorithms for prediction - the Preseq DS method based on rational function approximation (RFA), and the zero truncated negative binomial (ZTNB), where the a negative binomial distribution is fitted to the CU histogram and then used for prediction. The Preseq DS method is evaluated with 2 different parameterizations (MT = 2 and MT = 20), corresponding to different number of copies per UMI at which the CU histogram is truncated. In addition, we evaluated a selection method recommended by Deng et al (13), referred to as "best practice", in which Preseq DS (MT = 2) is chosen for genes where the CV of the counts per UMI is greater than one and ZTNB otherwise.

We downsampled 13 datasets to one tenth of the original amount of reads, and predicted up to the original gene expression. Fig. S6 A shows the $\log_2$ fold change (LFC) between corrected expression and ground truth for genes as a function of the number of UMIs available for the gene in the downsampled dataset. The LFC presented for each method was calculated as a loess fit of all genes from all datasets included in this study. Lin. scaled corresponds to the case where all genes are scaled equally (i.e. no advanced prediction is used). Fig. S6 B shows an analogous analysis, with the difference that the data has been CPM-normalized before the LFC is calculated. It is evident that CPM normalization improves the correction performance.

ZTNB and best practice are almost identical, suggesting that the ZTNB is chosen for most genes for the best practice case algorithm, and we see no benefit using the best practice method. In general, ZTNB has better performance for highly expressed genes, while Preseq DS is marginally better for the middle range genes, but the overall performance is similar. Evaluation for each of the 13 datasets individually is available in Fig. S8-S20. In general, the results are similar across datasets. The individual evaluation of datasets show that Preseq DS is more stable when CU histograms are truncated at 2 as compared to 20; the latter parametrization sometimes gives rise to outlier genes with larger prediction errors (e.g. Fig S19 D). Interestingly, the Preseq DS algorithm (MT = 20) performs best for predicting the total number of molecules in most cases (e.g. Fig. S20 A). We speculate that this method can take advantage of the high number of molecules, and that a negative binomial distribution may not be sufficient to describe the CU histogram of differently amplified pooled genes.
